# Bacterial outer-membrane polysaccharide export (OPX) proteins occupy three structural classes with selective β-barrel porin requirements for polymer secretion

**DOI:** 10.1101/2022.02.11.480155

**Authors:** Fares Saïdi, Utkarsha Mahanta, Adyasha Panda, Nicolas Y. Jolivet, Razieh Bitazar, Gavin John, Matthew Martinez, Abdelkader Mellouk, Charles Calmettes, Yi-Wei Chang, Gaurav Sharma, Salim T. Islam

**Author notes:** corresponding authors: Salim T. Islam,; Gaurav Sharma. co-1^st^ authors (Fares Saïdi, Utkarsha Mahanta).

## Abstract

Secretion of high-molecular-weight polysaccharides across the bacterial envelope is ubiquitous as it enhances prokaryotic survival in (a)biotic settings. Such polymers are often assembled by Wzx/Wzy- or ABC transporter-dependent schemes that implicate outer-membrane (OM) polysaccharide export (OPX) proteins in polymer translocation to the cell surface. In the social predatory bacterium *Myxococcus xanthus*, exopolysaccharide (EPS)-pathway WzaX, major spore coat (MASC)-pathway WzaS, and biosurfactant polysaccharide-pathway WzaB were herein found to be truncated OPX homologues of *Escherichia coli* Wza lacking OM-spanning α-helices. Comparative genomics across all bacteria, complemented with cryo-electron tomography cell- envelope analyses, revealed WzaX/S/B architecture to be the most common amongst three defined OPX-protein structural classes independent of periplasmic thickness. Fold-recognition and deep- learning analyses revealed the conserved *M. xanthus* proteins MXAN_7418/3226/1916 (encoded adjacent to WzaX/S/B) to be integral OM β-barrels, with structural homology to the poly-*N*-acetyl-D- glucosamine synthase-dependent pathway porin PgaA. Such porins were identified in bacteria near numerous genes for all three OPX-protein classes. Interior MXAN_7418/3226/1916 β-barrel electrostatics were found to match known properties of their associated polymers. With MXAN_3226 essential for MASC export, and MXAN_7418 absence shown herein to compromise EPS translocation, these data support a novel secretion paradigm for Wzx/Wzy-dependent pathways in which those containing an OPX component that cannot span the OM instead utilize a β-barrel porin to mediate polysaccharide transport across the OM.

## INTRODUCTION

Diverse bacteria associated with biotic and abiotic settings secrete high-molecular-weight (HMW) polysaccharides across the cell envelope to enhance their survival. Capsule polysaccharide (CPS) chains are tightly bound to the cell surface and form hydrated exclusionary barriers to molecule entry, whereas exopolysaccharide (EPS) polymers form a more loosely surface-associated glycocalyx around cells that contributes to biofilm matrix formation in aggregates (Whitfield *et al*., 2020). Certain HMW polysaccharides do not remain associated with the cell surface and are instead secreted to the extracellular milieu where they can influence bacterial physiology (Saïdi *et al*., 2021, Islam *et al*., 2020). In both monoderm (Gram-positive) and diderm (Gram-negative) bacteria, multiple secreted polymers often act in concert to modulate complex physiology (Pérez-Burgos & Søgaard-Andersen, 2020, Lavelle *et al*., 2021, Franklin *et al*., 2011).

*Myxococcus xanthus* is a Gram-negative bacterium that exhibits an intricate social multicellular lifestyle (Muñoz-Dorado *et al*., 2016). This motile soil bacterium (Faure *et al*., 2016, Islam & Mignot, 2015) is able to predate other bacteria (Seef *et al*., 2021) and saprophytically feed on the degradation products. Under nutrient deprivation, cells in a swarm biofilm form myxospore- filled fruiting bodies through a developmental program resulting in a differentiated cell community (Muñoz-Dorado *et al*., 2016). This complex lifecycle is modulated by the secretion of three known polysaccharides (Pérez-Burgos & Søgaard-Andersen, 2020). Cells constitutively produce EPS, a specific surface-associated polymer that forms a glycocalyx surrounding the cell body (Saïdi *et al*., 2021) and which constitutes the main matrix component in biofilms of this bacterium (Hu *et al*., 2013, Smaldone *et al*., 2014). A biosurfactant polysaccharide (BPS) is also synthesized, but is instead secreted to the extracellular milieu (Islam *et al*., 2020), where it functionally destabilizes the EPS glycocalyx, leading to a range of fundamental behavioural and surface-property changes at the single-cell level (Saïdi *et al*., 2021). The synergy between EPS and BPS secretion as well as the spatiospecific production patterns of the two polymers (Islam *et al*., 2020) also impacts the internal architecture of *M. xanthus* swarm biofilms, as well as their Type IV pilus (T4P)-dependent expansion due to impacts on T4P production, stability, and positioning (Saïdi *et al*., 2021). Finally, the <u>ma</u>jor <u>s</u>pore <u>c</u>oat (MASC) polymer is produced by spore-forming cells undergoing development, providing a protective coat to cover nascent myxospores and provide resistance to environmental stresses (Holkenbrink *et al*., 2014, Wartel *et al*., 2013).

Each *M. xanthus* polysaccharide is produced by a separate Wzx/Wzy-dependent assembly pathway (Islam *et al*., 2020, Pérez-Burgos *et al*., 2020, Holkenbrink *et al*., 2014), the components for which have the suffixes X (e<u>x</u>opolysaccharide), B (<u>b</u>iosurfactant), or S (<u>s</u>pore coat). Therein, cluster-specific glycosyltransferases synthesize polymer repeat units atop an undecaprenyl pyrophosphate (UndPP) lipid anchor at the cytoplasmic leaflet of the inner membrane (IM). UndPP-linked repeats are then translocated across the IM via the Wzx flippase (Islam *et al*., 2012, Islam *et al*., 2013a, Islam *et al*., 2010), followed by polymerization at the periplasmic leaflet of the IM by Wzy (Islam *et al*., 2011, Islam *et al*., 2013b) to modal lengths governed by the Wzc polysaccharide co-polymerase (PCP) in non-O-antigen systems (**Fig. 1A**) (Islam & Lam, 2014, Whitfield *et al*., 2020). In *M. xanthus*, WzcB possesses an integrated cytosolic bacterial tyrosine autokinase (BYK) domain (PCP- 2A class), whereas WzcX and WzcS do not (PCP-2B class); the EPS and MASC pathways thus encode a separate BYK (Wze) for association with the cognate PCP (Islam *et al*., 2020). The Wzb bacterial tyrosine phosphatase (BYP) protein in turn regulates the state of PCP-2A Wzc and PCP-2B- associated Wze phosphorylation (Mori *et al*., 2012). Wzb-mediated dephosphorylation of BYK domains has been proposed to drive Wzc octamerization, in turn affecting Wzy-mediated polymerization and interaction with the outer-membrane polysaccharide export (OPX) Wza translocon needed for polymer transport through the periplasm and across the outer membrane (OM) (Yang *et al*., 2021) (**Fig. 1A**). Such pathways are one of the most widespread bacterial assembly schemes for HMW polysaccharides, responsible for synthesizing diverse products such as Group 1 CPS and Group 4 CPS (i.e. “O antigen” capsule) as well as colanic acid polymers in enterobacteria (Sande & Whitfield, 2021), in addition to holdfast polysaccharide in *Caulobacter crescentus*, (Toh *et al*., 2008), and xanthan in *Xanthomonas campestris* (Becker, 2015). In Group 1 CPS systems, the 18- stranded integral OM β-barrel Wzi (internally occluded by an α-helical plug domain) is also important as it displays lectin-like characteristics implicated in capsule structure organization (Bushell *et al*., 2013).

**Figure 1.**
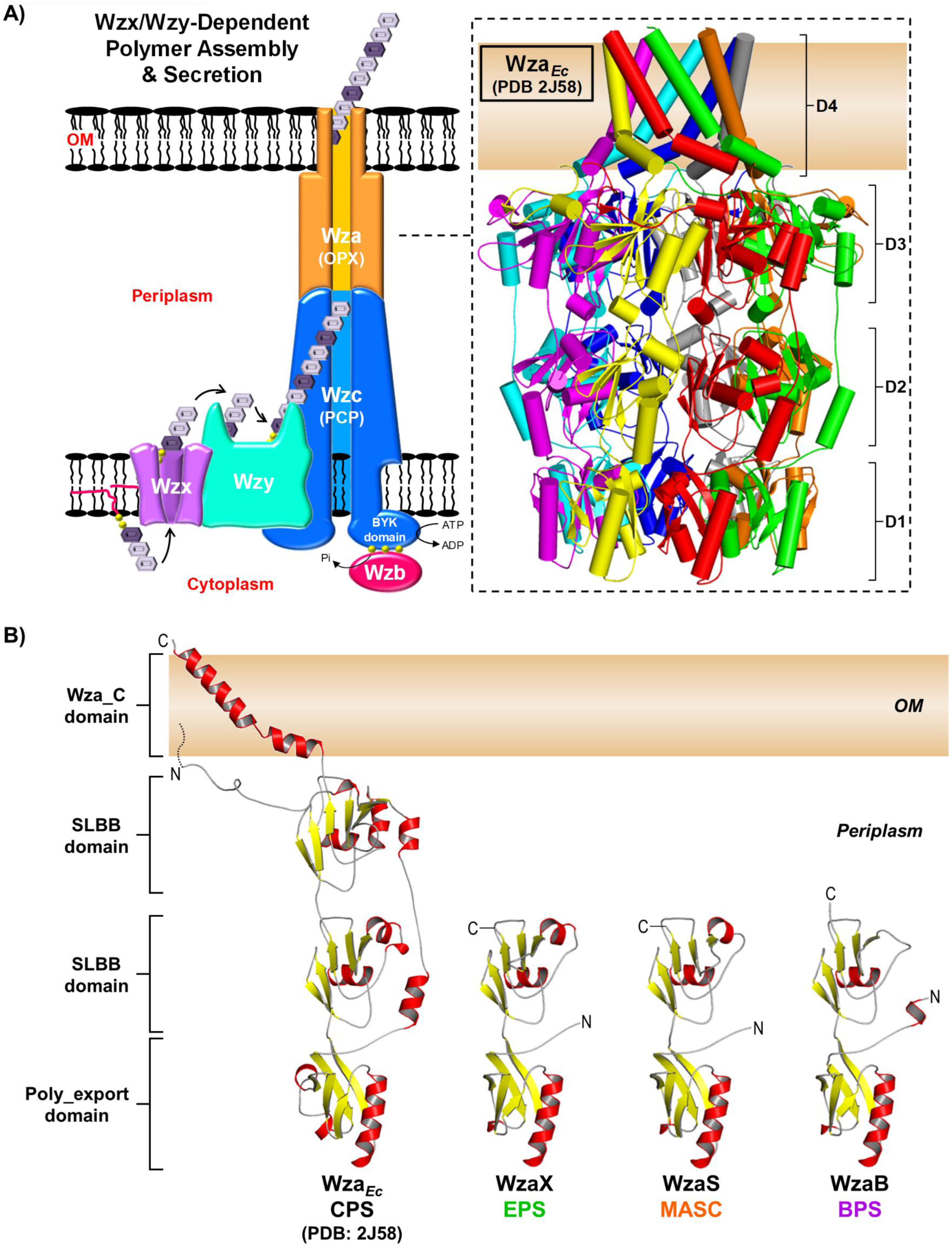
Wzx/Wzy-dependent polysaccharide assembly-and-secretion. **A)** Pathway schematic. *Inset:* The Wza*_Ec_* X-ray crystal structure octamer (PDB: 2J58) has been differentially coloured to highlight the position of each chain in the structure. The D1 (Poly_export), D2 & D3 (both SLBB), and D4 (Wza_C) domains have been indicated, with smooth loops. **B)** Tertiary structure models of *M. xanthus* EPS-pathway WzaX (aa 51–212), MASC-pathway WzaS (aa 32–190), and BPS-pathway WzaB (aa 38–202) based on the Wza*_Ec_* structure (aa 22–376, depicted with a N-terminal lipid anchor). Structures are displayed with smooth loops, highlighted β-sheets (*yellow*) and α-helices (*red*), with the various N- and C- termini indicated.

Secreted polymers can also be synthesized by an ABC transporter-dependent scheme in which UndPP-linked polymers are built by sugar unit addition at the cytoplasmic leaflet of the IM, with the polymer generated entirely in the cytoplasm. Subsequent ATP hydrolysis by the transporter drives polymer transport across the IM, after which PCP and OPX proteins are needed for the secretion of polymer through the periplasm and across the OM. These biosynthesis pathways are implicated in the secretion of polysialic acid Group 2 CPS polymers from pathogenic extraintestinal *E. coli*, as well as similar structures in *Neisseria meningitidis* and *Haemophilus influenza* (Willis & Whitfield, 2013).

Alternatively, HMW polymers such as alginate in *Pseudomonas aeruginosa*, cellulose in *Salmonella enterica*, and poly-*N*-acetyl-D-glucosamine (PNAG) in *Acinetobacter baumanii* are produced via a synthase-dependent scheme in which the addition of a monosaccharide in the cytoplasm by an integral IM synthase results in export of the polymer by a similar amount from the cell surface. Polymer transport through the periplasm is mediated by a protein scaffold containing TPR repeats followed by translocation across the OM through an integral OM β-barrel porin structure (Whitney & Howell, 2013). While similarities exist between Wzx/Wzy- and ABC transporter- dependent pathways (e.g. the presence of PCP and OPX proteins), no such schematic crossover with proteins from synthase-dependent pathways has been identified.

In both Wzx/Wzy- and ABC transporter-dependent pathways, OPX-family proteins are portrayed as forming an oligomeric channel of contiguous domains to allow polymer secretion through the periplasm and across the OM (Whitfield *et al*., 2020). All OPX proteins share a conserved periplasmic N-terminal Poly_export domain (Pfam: PF02563, also called a PES [polysaccharide export sequence] motif), followed by at least one copy of a soluble ligand-binding β- grasp (SLBB) domain (Pfam: PF10531) (Sande *et al*., 2019, Cuthbertson *et al*., 2009), which is predicted to interact with the sugar polymer in the periplasm (Burroughs *et al*., 2007). OPX protein domain architecture diverges at this point. In the prototypic OPX Wza from *E. coli* group 1 CPS (Wza*Ec*) — which is the only OPX protein with a solved 3D structure — after one Poly_export (D1) and two SLBB domains (D2 & D3), the protein contains a Wza_C domain (Pfam: PF18412) at its C- terminus, which forms a 35-residue amphipathic α-helical tract (D4) that crosses the OM (Dong *et al*., 2006) (**Fig. 1A**). Elucidation of the Wza*Ec* X-ray crystal structure (PDB: 2J58) was revolutionary as it represented the first instance of such a fold in an integral OM protein. As part of the Wza*Ec* oligomer, 8 copies of the α-helical Wza_C domain were shown to form a pore-like structure, through which it is proposed that secreted polysaccharides exit the cell (Dong *et al*., 2006, Nickerson *et al*., 2014) (**Fig. 1A**). Instead of a classical Wza_C domain, OPX proteins from ABC transporter-dependent Group 2 CPS pathways usually contain a C-terminal Caps_synth_GfcC (Pfam: PF06251, formerly DUF1017) module (Cuthbertson *et al*., 2009), structurally similar to the stand- alone GfcC protein (PDB: 3P42) from Group 4 Wzx/Wzy-dependent CPS pathways (Sande *et al*., 2019). GfcC contains domains comparable to D2 and D3 from Wza*Ec*, as well as a D4-like amphipathic α-helix; however, the GfcC D4-like helix spanning the final 21 residues of the protein is 40% shorter than Wza*Ec* D4, bent at both ends, and structurally locked (Sathiyamoorthy *et al*., 2011). This overall OPX architecture is typified by the Group 2 CPS pathway protein KpsD from *E. coli* (KpsD*Ec*) (Sande *et al*., 2019). Though it is uncertain if the C-terminal domain of KpsD*Ec* is able to span the OM, KpsD*Ec* epitopes have previously been detected at the cell surface via anti-KpsD*Ec* antibody labelling (McNulty *et al*., 2006). Numerous other annotated OPX proteins have been shown to either (i) contain considerable-yet-uncharacterized protein sequences following their most C-terminal identified domain, or (ii) be considerably shorter than either Wza*Ec* or KpsD*Ec*, with architecture beyond the Poly_export and SLBB domains largely absent (Cuthbertson *et al*., 2009). Ultimately, for OPX proteins that lack a canonical OM-spanning Wza_C domain, the manner by which the respective secreted polymers traverse the asymmetric OM bilayer remains a fundamental and pertinent question that has yet to be resolved.

Herein, we reveal that the WzaX, WzaB, and WzaS OPX proteins for the respective *M. xanthus* EPS, BPS, and MASC pathways contain typical N-terminal “Poly_export–SLBB” architecture but lack a C-terminal OM-spanning Wza_C domain. Comparative genomics analyses reveal this architecture to be the most common amongst three distinct structural classes of OPX proteins across all bacteria. However, in the *M. xanthus* EPS, BPS, and MASC biosynthetic clusters, a conserved β-barrel protein (MXAN_7418/MXAN_1916/MXAN_3226) is encoded immediately adjacent to the genes for WzaX/WzaB/WzaS (respectively). Fold-recognition and deep-learning analyses reveal these adjacently-encoded proteins to be 18-stranded integral OM β-barrels with structural homology to the barrel domain of the porin PgaA, required for PNAG secretion across the OM by synthase-dependent pathways. In turn, PgaA-like β-barrel proteins are shown to be encoded near numerous genes representing all three OPX structural classes in diverse Gram-negative bacteria. The interior electrostatics of the *M. xanthus* β-barrels match known properties of their associated polymers, and deletion of the MXAN_7418 β-barrel is shown to compromise EPS secretion.

Together with the known requirement for the MXAN_3226 β-barrel for MASC secretion (Holkenbrink *et al*., 2014), these data support a novel secretion paradigm for Wzx/Wzy-dependent pathways in which those containing an OPX component that cannot span the OM instead utilize a β- barrel porin to mediate translocation of HMW polymers across the OM.

## RESULTS

### The *M. xanthus* OPX proteins WzaX, WzaS, and WzaB lack an OM-spanning α-helix domain

Each of the WzaX/S/B OPX proteins is essential for the secretion of its respective EPS/MASC/BPS polymer in *M. xanthus* (Islam *et al*., 2020, Holkenbrink *et al*., 2014). However, as each of WzaX, WzaS, and WzaB has a considerably shorter amino acid sequence than that of Wza*Ec* (44, 50, and 46% smaller, respectively), we sought to better understand the structural implications of this difference. Fold-recognition analysis of each protein revealed 100%, 100%, and 99.9% probability matches (respectively) to the N-terminal half of the high-resolution Wza*Ec* X-ray crystal structure. However, tertiary structure modelling of WzaX, WzaS, and WzaB against the Wza*Ec* 3D structure revealed WzaX, WzaS, and WzaB to be missing the 2^nd^ SLBB domain (i.e. D3), and more importantly the crucial OM-spanning α-helical Wza_C domain (i.e. D4), of Wza*Ec* (**Fig. 1B**). The absence of such an OM-spanning domain is consistent with the lack of WzaX/S/B detection in proteomic analyses of OM vesicle (OMV) and biotinylated surface-protein samples (Kahnt *et al*., 2010), despite the constitutive expression of the *wzaX/S/B* genes throughout the *M. xanthus* lifecycle (Muñoz-Dorado *et al*., 2019, Sharma *et al*., 2021).

### OPX proteins constitute three distinct structural classes

To determine if the absence of the Wza_C domain was an aberration confined to the subset of OPX proteins from *M. xanthus* under study, we first performed a comparative genomics analysis using profile-based homology searches across three different datasets: (i) 61 myxobacterial genomes (MYXO) (**Supplementary Table S1**), (ii) 3662 representative genomes (REP) (**Supplementary Table S2**), and (iii) the non-redundant (NR) NCBI database (371 327 556 proteins at 100% identity as of June 10, 2021) (**Supplementary Table S3**), to identify encoded OPX proteins, using PF02563 [Poly_export], PF10531 [SLBB], PF18412 [Wza_C], and PF06251 [Caps_synth_GfcC; used here as “GfcC”] as our query domains. These profile-based analyses identified diverse putative OPX homologues that we divided into three distinct classes according to their domain architecture (**Fig. 2A**). The first set of OPX proteins was found to contain Poly_export–SLBB(1–14) architecture ending with a C-terminal OM-spanning Wza_C domain, similar to Wza*Ec*, and was assigned the designation “Class 1” (**Fig. 2A**). A second set of OPX proteins was found to possess Poly_export– SLBB(1–6)–GfcC architecture similar to KpsD*Ec*, ending with or without a C-terminal OM-spanning Wza_C domain, and was assigned the designation “Class 2”. However, most OPX proteins were found to contain only Poly_export–SLBB(1–7) architecture lacking either a Wza_C or GfcC domain; these hits were designated “Class 3”; however, many of these initial hits were found to contain additional amino acids that may have remained uncharacterized following sequence-based domain detection. Therefore, to probe these partially-characterized hits in more detail, we subjected all identified OPX proteins to fold-recognition analysis using HHpred to identify matches with more remote sequence homology but conserved structural properties. These analyses resulted in reclassification of several original Class 3 hits to either Class 1 or Class 2

**Figure 2.**
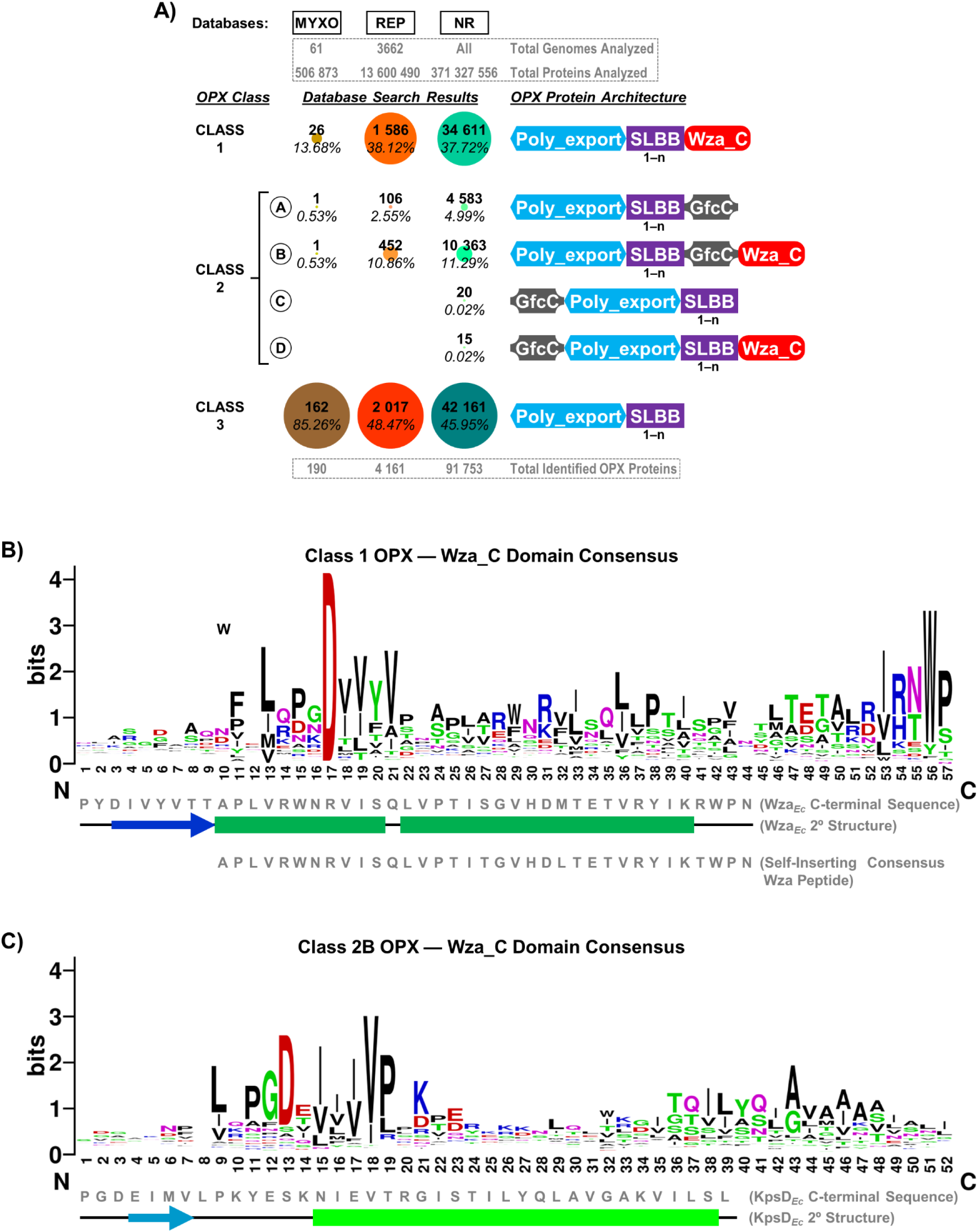
Structural diversity of OPX proteins. **A)** Domain organization and abundance of bacterial OPX protein classes identified in the myxobacterial (MYXO, 61 genomes, 506 873 proteins analyzed), representative (REP, 3662 genomes, 13 600 490 proteins analyzed), and NCBI non- redundant (NR, 371 327 556 proteins analyzed) databases. The Poly_export (PF02563), SLBB (PF10531), Wza_C (PF18412), and Caps_synth_GfcC (PF06251) Pfam domains were used to query the various databases, followed by fold-recognition analysis using HHpred against the 3D X- ray crystal structures of Wza*_Ec_* (PDB: 2J58) and GfcC (PDB: 3P42). The number of repeated copies is indicated under each domain depiction. The number of OPX hits (*bold*) for a specific class is indicated as well as the proportion of hits from each database (*italics*) represented by the hits. **B)** Sequence logo of the consensus amino acids constituting the OM-spanning α-helix based on a multiple-sequence alignment of 1 586 Class 1 OPX proteins. The region of sequence alignment with Wza*_Ec_* is indicated and depicted with the associated secondary structure from the Wza*_Ec_* X-ray crystal structure (PDB: 2J58) (Dong *et al*., 2006). Also depicted is the region of sequence alignment with a previously-published optimized Wza_C synthetic peptide (based on 94 close Wza*_Ec_*-related homologues) capable of spontaneously inserting into lipid bilayers and self-assembling into stable α- barrel pores (Mahendran *et al*., 2017). **C)** Sequence logo of the consensus amino acids constituting the putative OM-spanning α-helix based on a multiple-sequence alignment of 452 Class 2B OPX proteins. The region of sequence alignment with KpsD*_Ec_* is indicated, along with the predicted KpsD*_Ec_* secondary structure. The position of observed (*dark-colored*) and predicted (*light- colored*) α-helices (*boxes*) and β-strands (*arrows*) have been indicated.

Incidentally for Class-1 OPX proteins, while the secondary structure was conserved, considerable sequence variation was detected within certain regions of the putative OM-spanning Wza_C domains, with this domain extending up to 48 residues in length (compared to 35 residues in Wza*Ec*) (**Fig. 2B**). As per the MYXO/REP/NR databases, 13.7/38.1/37.7% of OPX proteins possess Class 1 Wza*Ec*-like organization with a putative OM-spanning C-terminal α-helix (**Fig. 2A, Supplementary Tables S1, S2, S3**). Class 1 OPX proteins were found to have a median length of 378 amino acids and most (1233/1586, ∼78%) were predicted to be lipoproteins with Sec/SPII signal sequences.

These proteins were largely confined to the phylum Proteobacteria (837/1632 genomes; ∼51%) with a predominance in classes Gammaproteobacteria (430/760 genomes; ∼56%), Alphaproteobacteria (232/410 genomes; ∼57%), and Betaproteobacteria (133/274 genomes; ∼49%), and representation also in phyla Bacteroidetes (67/283 genomes; ∼24%), Planctomycetes (56/62 genomes; ∼90%), and Cyanobacteria (46/55 genomes; ∼84%) (**Supplementary Table S2**). Based on species-level PSORTdb classification, the REP database contains 698 Gram-positive and 1381 Gram-negative organisms. Our analysis revealed that Class 1 OPX proteins are encoded by many Gram-negative bacteria (639/1381 genomes, ∼46%), whereas these proteins were completely absent in Gram-positive species (**Supplementary Table S2**).

Our MYXO/REP/NR database comparative genomic analysis revealed that Class 2 OPX proteins can be further divided into four subclasses. Proteins belonging to Class 2A contain Poly_export–SLBB(1–n)–GfcC architecture, whereas those assigned to Class 2B possess Poly_export– SLBB(1–n)–GfcC–Wza_C architecture ending with an OM-spanning α-helical domain. Classes 2C and 2D are variations of Classes 2A and 2B (respectively) where the Poly_export domain is preceded by a GfcC domain; however, only 20 Class 2C and 15 Class 2D proteins were identified across the entire NR database.

Class 2A OPX proteins constitute 0.5/2.6/5.0% of all OPX proteins identified in the MYXO/REP/NR databases, with a median length of 605 amino acids. These proteins were found to be encoded mainly in Proteobacteria (77/1632 genomes, ∼5%), Bacteroidetes (9/283 genomes, ∼3%), and Acidobacteria (3/14 genomes, ∼21%). Taxonomy orders Alteromonadales (14/83 genomes, ∼17%), Campylobacterales (10/88 genomes, ∼11%), Burkholderiales (9/175 genomes, ∼5%), and Oceanospirillales (9/43 genomes, ∼21%) display the maximum representation for Class 2A. In addition, Class 2A OPX proteins are encoded in only ∼5% (66/1381 genomes) of Gram-negative bacteria and are completely absent among Gram-positive species (**Supplementary Table S4**).

Class 2B OPX proteins represent 0.5/10.9/11.3% of all OPX proteins identified in the MYXO/REP/NR databases (**Fig. 2A, Supplementary Table S1, S2, S3**). The median length of Class 2B proteins was found to be 824 amino acids, with most (375/452, ∼83%) found to possess standard Sec/SPI secretory signal peptides. These proteins were largely encoded by Proteobacteria (226/1632 genomes, ∼14%), Bacteroidetes (107/283 genomes, ∼38%) and Cyanobacteria (14/55 genomes, ∼25%). At the level of Order, Alteromonadales (66/83 genomes, ∼80%), Bacteroidales (38/52 genomes, ∼73%), Cytophagales (23/50 genomes, 46%), and Vibrionales (20/46 genomes, ∼43%) were found to contain the most organisms encoding Class 2B OPX architecture. Class 2B OPX proteins have representation in only ∼18% (243/1381 genomes) of Gram-negative organisms and are absent in Gram-positive bacteria (**Supplementary Table S4**).

Class 2B architecture is typified by KpsD*Ec*. Consistent with a previous report (Sande *et al*., 2019), fold-recognition analysis of KpsD*Ec* revealed that most of its N-terminus is structurally homologous to Wza*Ec*, while the bulk of its C-terminus is a structural match to the standalone GfcC protein (**Supplementary Fig. S1A**). However, the extreme C-terminus of KpsD*Ec* — i.e. the portion of KpsD*Ec* surpassing the end of structural homology with the GfcC D4 α-helix — was found to have considerable structural homology with the most C-terminal region of Wza*Ec*, including a 25- residue tract with α-helical propensity matched to the OM-spanning α-helical tract of Wza*Ec* (**Supplementary Fig. S1B,C**). A similarly-extended C-terminal α-helix was found throughout the Class 2B OPX hits identified herein, with considerable variation in certain regions of its sequence, and extending to 38 residues (compared to 25 residues in KpsD*Ec*) (**Fig. 2C**). This observation supports the notion that a part of KpsD*Ec* (and by extension Class 2B OPX proteins) may indeed be able to span the OM and access the cell surface.

Finally, Class 3 OPX proteins with Poly_export–SLBB(1–n) architecture (but no appreciable peptide sequence following their respective C-terminal-most SLBB domain), with a median length of 256 amino acids, represent a plurality (∼85/49/46%) of OPX proteins identified across the MYXO/REP/NR databases (**Fig. 2A, Supplementary Table S1, S2, S3**). Almost 50% are predicted lipoproteins (Sec/SPII signal sequences) while ∼30% are secreted proteins (Sec/SPI signal sequences). Such OPX proteins (i.e. those lacking a Wza_C or GfcC) domain were found across multiple bacterial classes such as Alphaproteobacteria (274/410 genomes, ∼67%), Gammaproteobacteria (247/760 genomes, ∼33%), Betaproteobacteria (100/274 genomes, ∼37%), Flavobacteria (116/133 genomes, ∼87%), and Deltaproteobacteria (66/82 genomes, ∼80%).

Expectedly, our analysis detected Class 3 OPX proteins in Gram-negative bacteria (600/1382 genomes, ∼43%), but also intriguingly in several Gram-positive organisms (52/699 genomes, ∼7%) (**Supplementary Table S4**). Of note, the proportions of each Class of OPX protein detected in the REP database were highly reflective of those found in the NR database (**Fig. 2A**), reinforcing the utility and applicability of the REP database.

Within the MXYO dataset, we identified 26 Class 1, two Class 2, and 162 Class 3 OPX proteins (**Fig. 2A**, **Supplementary Table S1**). Class 3 OPX proteins were encoded in all 61 myxobacterial organisms, without any exception, in the range of 1-4 proteins. Class 1 OPX proteins were encoded by several members of the suborder Cystobacterineae such as *Anaeromyxobacter*, *Cystobacter*, *Corallococcus*, *Pyridicoccus*, and *Simulacricoccus*. However, Class 1 OPX proteins were not represented within any species of the genus *Myxococcus*. Class 2 OPX proteins were only present in two myxobacteria, namely *Sandaracinus amylolyticus* (Class 2A OPX) and *Haliangium ochraceum* (Class 2B OPX). All proteins in the MYXO dataset belonging to the three OPX classes possess similar median lengths to those described for all OPX proteins in the REP database (i.e. MYXO Class 1, 373 amino acids; MYXO Class 2, 550 amino acids; MYXO Class 3, 205 amino acids).

### Molecular phylogenetics suggests the coevolution of three OPX protein Classes

Taken together, these analyses have identified three principal structural classes of OPX proteins, namely: (i) Class 1 (i.e. Wza*Ec*-like with a canonical OM-spanning α-helix), (ii) Class 2 (i.e. KpsD*Ec*-like with a potentially OM-spanning α-helix at the terminus of the GfcC domain), and (iii) the majority Class 3 (i.e. those with structural homology to the Wza*Ec* N-terminus, but with no discernible OM-spanning domain). By extension, WzaX/S/B from *M. xanthus*(**Fig. 1B**) are thus Class 3 OPX proteins that share the domain architecture of most other OPX proteins encoded by bacteria (**Fig. 2A**). Given that all OPX proteins among the three different classes have a conserved “Poly_export” domain, this can be utilized as a phylogenetic marker. Therefore, based on the hmmscan results, we extracted the location of the “Poly_export” domain from REP dataset hits, aligned those sequences using MUSCLE, and generated a maximum-likelihood phylogeny. The generated phylogeny (**Fig. 3**) revealed that Class 1 and Class 3 OPX proteins are interspersed with each other in all taxonomic clades, suggesting that proteins from these two classes have co-evolved by losing or gaining the Wza_C segment in closely-related organisms. However, Class 2A and Class 2B are both present in nearby sister clades and away from Class 1 and Class 3. This denotes that Class 2 OPX proteins, similar to proteins from Class 1 and Class 3, are highly similar to each other and have coevolved by losing or gaining their respective Wza_C segments.

**Figure 3.**
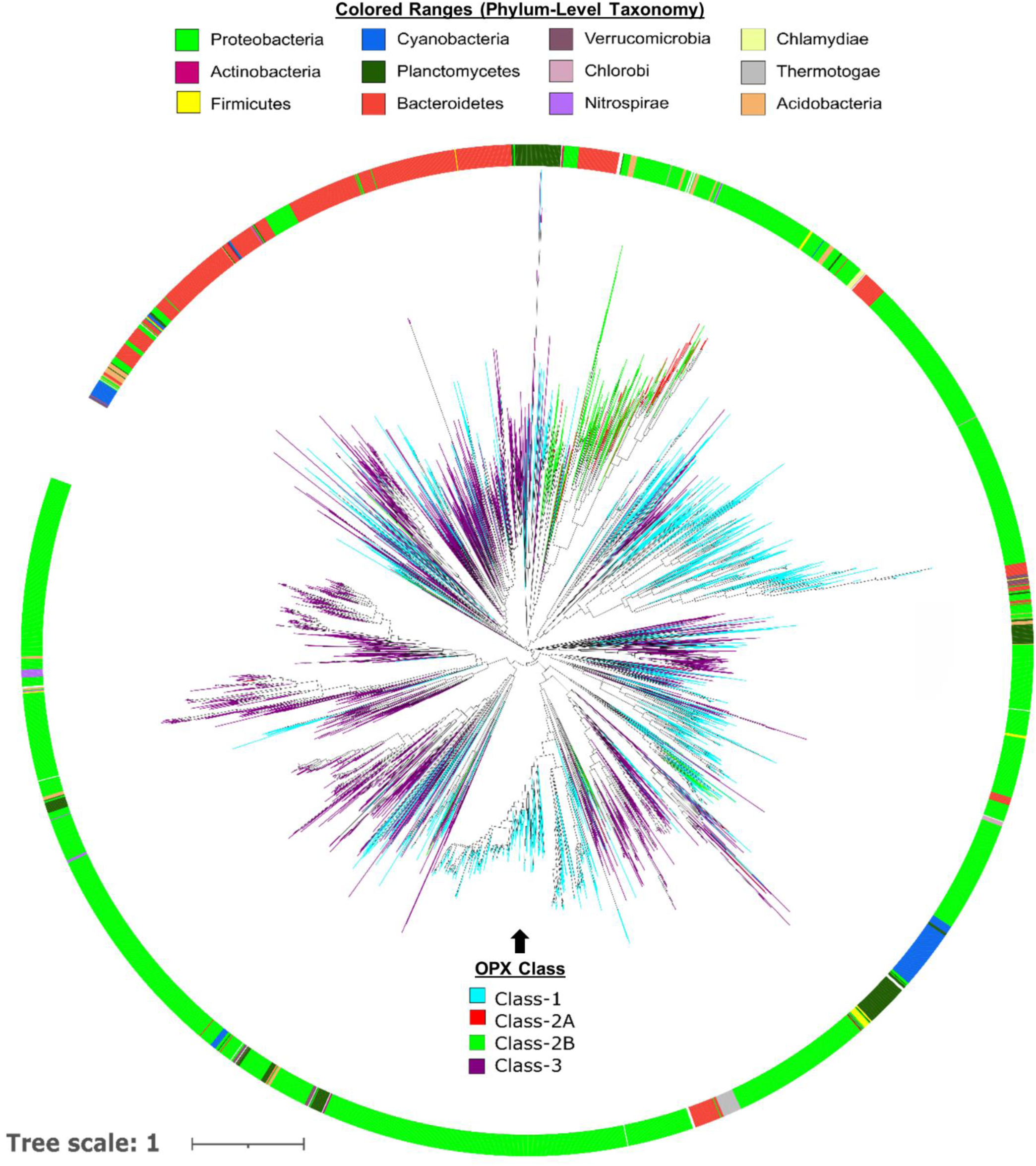
Phylogenetic tree of OPX proteins. Sequence alignment of all “Poly_export” domains as identified in 4 161 OPX proteins in the REP dataset was used to generate a maximum-likelihood phylogenetic tree. The classes of OPX proteins (*inner tree*) and their respective phyla (*outer ring*) have been coloured accordingly for effective visualization.

### OPX protein lengths and classes in Gram-negative bacteria are not linked to periplasmic distance

In *E. coli*, the integral OM Class 1 Wza*Ec* OPX protein is proposed to form a complex with the integral IM Wzc PCP protein that creates a contiguous periplasm-spanning channel for polymer export (Collins *et al*., 2007) (**Fig. 1A**). To gain an understanding of the relationship between the subcellular architecture of *M. xanthus* and the role of the WzaX, WzaS, and WzaB Class 3 OPX proteins, we therefore compared the sizes of various cellular compartments and structures from cryo- electron tomography projections of the *M. xanthus* envelope. This revealed the *M. xanthus* OM to have an average thickness of 69.8 ± 1.8 Å, compared to the average thickness of 62 ± 1.6 Å for its IM, with a mean inter-membrane periplasmic thickness of 327 ± 28.4 Å (**Fig. 4A**).

**Figure 4.**
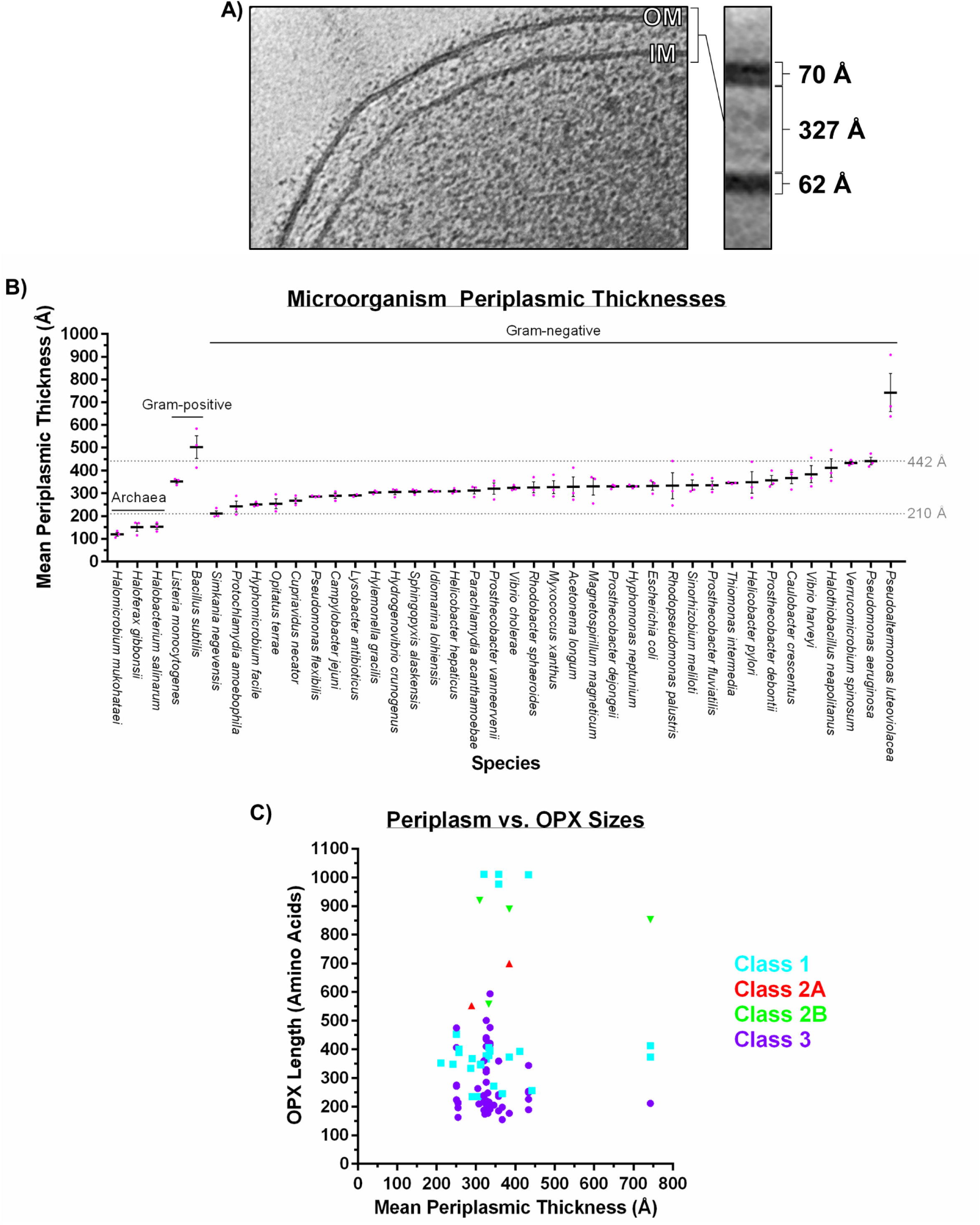
Cell envelope ultrastructure in Gram-negative bacteria. **A)** Cryo-electron microscopy tomogram slice of a *M. xanthus* cell, showing the IM, OM, and intervening periplasmic space and their respective measured thicknesses. **B)** Comparison of means (*black bar*) for periplasmic distances in 40 microbial species (± SEM). Individual replicate measurements are indicated (*red dots*). Data from organisms with increasing mean periplasmic thickness values have been depicted from left to right, grouped according to Gram-negative, Gram-positive, or Archaea organism designation. **C)** Scatterplot of mean periplasmic thickness values plotted against the length of each OPX protein from the same organism. Data points have been coloured according to the Class of OPX protein assigned herein. No correlation was detected between periplasmic thickness and (i) overall OPX protein length (Pearson coefficient: 0.1450; Spearman coefficient: 0.05884, calculated over 92 data pairs), (ii) Class 1 OPX hits (Pearson coefficient: 0.07250; Spearman coefficient: 0.2318; calculated over 33 data pairs), or (iii) Class 3 OPX hits (Pearson coefficient: -0.1127; Spearman coefficient: -0.1242; calculated over 53 data pairs).

Given the enrichment of Class 1 OPX proteins in certain bacterial genera and different median OPX sizes for each OPX protein class (**Supplementary Table S4**), we examined whether the specific size of an OPX protein identified in a given bacterium was associated with the thickness of the periplasm in that organism. We first measured the distance between the IM and OM at lateral positions in cryo-electron tomography projections of cells from an additional 34 species of Gram-negative bacteria. For reference, we also analyzed IM–peptidoglycan and IM–S-layer thickness in several Gram-positive bacteria and Archaea (respectively) (**Fig. 4B**). This analysis revealed a range of Gram-negative periplasmic thicknesses confined between the lower and upper thresholds of 210 and 442 Å (respectively), with the lone exception being *Campylobacter jejuni*, displaying a mean periplasmic thickness of 743 Å (**Fig. 4B**). For any of these species in which OPX proteins were herein identified (**Supplementary Tables S1, S2, S3**), we next compared the average measured periplasmic thickness with the length of the OPX protein(s) in each system. However, no overall correlation between the two variables was detected across all OPX proteins in this analysis, nor specifically within Class 1 or Class 3 OPX hits (**Fig. 4C**).

### WzaX, WzaS, and WzaB are genomically paired with 18-stranded β-barrel proteins

Given the lack of identifiable OM-spanning domains in WzaX/S/B (**Fig. 1B**), we sought to identify candidate proteins that could permit export of synthesized EPS, MASC, and BPS polymers across the *M. xanthus* OM. Through our previous analyses of the EPS, MASC, and BPS biosynthesis clusters, we demonstrated that WzaX (MXAN_7417), WzaS (MXAN_3225), and WzaB (MXAN_1916) were encoded immediately adjacent to MXAN_7418, MXAN_3226, and MXAN_1916 (respectively), with this synteny conserved for the majority (115/162, ∼71%) of Class 3 OPX proteins in myxobacterial genomes (Islam *et al*., 2020), supporting the notion that the latter three proteins are important for each respective polymer synthesis pathway. To analyze the specific structural potential for each protein, MXAN_7418, 3226, and 1916 were first subjected to evolutionary coupling analysis, revealing the predicted presence of 18 principal β-strands for MXAN_7418, MXAN_3226, and MXAN_1916 (**Supplementary Figs. 2A,3A,4A**).

Fold-recognition analyses revealed structural homology of the C-terminal 76-86% of MXAN_7418/3226/1916 to the complete C-terminal integral OM β-barrel module (residues 513-807) of PgaA (PgaAβb, PDB: 4Y25) (Wang *et al*., 2016), at 98.9%, 99.4%, and 99.2% probability (respectively). PgaA is the OM porin responsible for secretion of PNAG polymer (which is heavily implicated in biofilm integrity) in Gram-negative bacteria; it contains multiple periplasmic tetratricopeptide repeats at its N-terminus, followed by a 16-stranded integral OM β-barrel domain closed by four extracellular loops (Wang *et al*., 2016). Intriguingly, PNAG is produced by a synthase-dependent pathway (Whitney & Howell, 2013). In each of MXAN_7418/3226/1916, the PgaAβb-like module is extended by two integral OM β-strands at the N-terminus of each protein, suggesting that MXAN_7418, 3226, and 1916 do indeed have the propensity to form 18-stranded β- barrels (**Supplementary Figs. 2B,3B,4B**).

For the 162 Class 3 OPX proteins identified across 61 myxobacterial genomes (with Poly_export—SLBB1-2 architecture), most (115/162, ∼71%) were found to be encoded near an extended PgaAβb-like protein, whereas the 26 Class 1 (with Poly_export—SLBB1-2—Wza_C organization) and two Class 2 OPX proteins were not encoded near any such β-barrel protein (**Supplementary Table S1**). We again expanded our analysis beyond *M. xanthus* to determine whether the presence of a β-barrel porin was a common occurrence in pathways containing an OPX protein. Intriguingly in *E. coli*, the *gfcABCDE-etp-etk* cluster needed for Wzx/Wzy-dependent Group 4 CPS production, encodes the OPX protein GfcE (formerly YccZ/Wza22min) as well as the protein GfcD (formerly YmcA) (Peleg *et al*., 2005). The separate *yjbEFGH* (paralogous to *gfcABCD*) operon implicated in polysaccharide secretion encodes the GfcD-like protein YjbH (Ferrières *et al*., 2007). Both GfcD and YjbH are β-barrel proteins, which were recently identified to be part of a novel class of OM proteins (lacking published structures) with two separate β-barrels predicted to be formed by the same polypeptide chain (Solan *et al*., 2021). Herein, fold-recognition analysis revealed the N-terminal halves to be matches to the β-barrel amyloid transporter FapF from *Pseudomonas*, whereas the C-terminal halves (GfcDCterβb and YjbHCterβb, respectively) possessed structural homology to the PNAG PgaAβb module described above (**Supplementary Fig. 5A**).

This double-barrel arrangement was supported by AlphaFold2-generated deep-learning structure models for both full-length GfcD and YjbH (**Supplementary Fig. 5B**), with the larger barrel portions displaying considerable sequence homology to PgaAβb (**Supplementary Fig. 5C**).

To probe for the presence of similar β-barrels encoded near other OPX proteins, we used sequence homology searches (BLAST and HMMER) to examine the genomic context (up to 10 genes upstream and downstream) of the various OPX proteins we identified in the REP dataset, beginning with the β-barrel sequences of MXAN_7418, MXAN_3226, and MXAN_1916. Given the homology of the above proteins to PgaA, we added the PgaAβb, GfcDCterβb, and YjbHCterβb sequences as well. In addition, the β-barrel sequences of BcsC (BcsCβb, from PDB: 6TZK) (Acheson *et al*., 2019) and AlgE (from PDB: 4AFK) (Tan *et al*., 2014) were also included, given their porin functions in the synthase-dependent cellulose and alginate production pathways, respectively (Whitney & Howell, 2013). Finally, we also included the sequence of Wzi (PDB: 2YNK) lacking the plug domain (Wziβb), as this is an 18-stranded β-barrel known to be linked with polysaccharide biosynthesis clusters (Bushell *et al*., 2013).

Altogether, this analysis detected 365 β-barrel query homologues encoded near 344 OPX proteins of all three classes (**Table 1**). Of the 156 matches to the Wziβb query, 128 were of comparable size and conserved alignment to full-length Wzi (including the α-helical plug domain), consistent with these hits not being likely porin candidates. However, 28 homologues were a match to only Wziβb (i.e. no plug domain), reinforcing their candidacies as trans-OM export β-barrels; intriguingly, HHpred analysis of these 28 hits also revealed many with strong similarity to DUF6029, ascribing a potential polysaccharide secretion role to this heretofore uncharacterized protein domain. The PgaAβb query detected 12 homologues, 2 of which were uniquely-matched to only the β-barrel domain, whereas the remaining 10 aligned with full-length PgaA, consistent with these latter hits possessing the numerous N-terminal TPR domains as well as the C-terminal PgaAβb porin domain. These results were similar for BcsCβb, i.e. that of the 5 homologues detected near OPX proteins, each was a match to full-length BcsC indicating a conservation of N-terminal TPR and C-terminal porin domains (with no matches confined to BcsCβb detected). Fifteen homologues to synthase-dependent pathway AlgE were also detected. Profile matches to GfcDCterβb and YjbHCterβb queries (68 and 84, respectively) were largely matched via length and conserved alignment to full-length GfcD and YjbH, suggesting that these homologues contain the N-terminal FapF-like β-barrel domain as well as the C-terminal PgaAβb-like polymer-secretion domains; only two GfcDCterβb and three YjbHCterβb hits were identified with homology to only the C-terminal barrels of each protein. However, the GfcDCterβb homologue detected in *Aquifex aeolicus* (WP_164930611.1), as well as the hits to YjbHCterβb detected in *A. aeolicus* (WP_010880290.1) and *Sulfitobacter pseudonitzschiae* (WP_174861591.1), were only matched to the respective polysaccharide-secretion modules, indicating the potential for stand-alone export of sugar chains in these systems. Finally, homologues to MXAN_7418, MXAN_3226, and MXAN_1916 were only found to be encoded in myxobacterial genomes. Importantly, the presence of these extended PgaAβb-like 18-stranded β-barrels in myxobacteria was linked to nearby Class 3 OPX proteins. However, unlike OPX hits from the MYXO database, only 77/2017 Class 3 OPX proteins from the REP dataset were identified to be encoded near MXAN_7418/3226/1916/ PgaAβb/GfcDCterβb/YjbHCterβb/BcsCβb/AlgE/Wziβb stand-alone porin homologues.

**TABLE 1.**
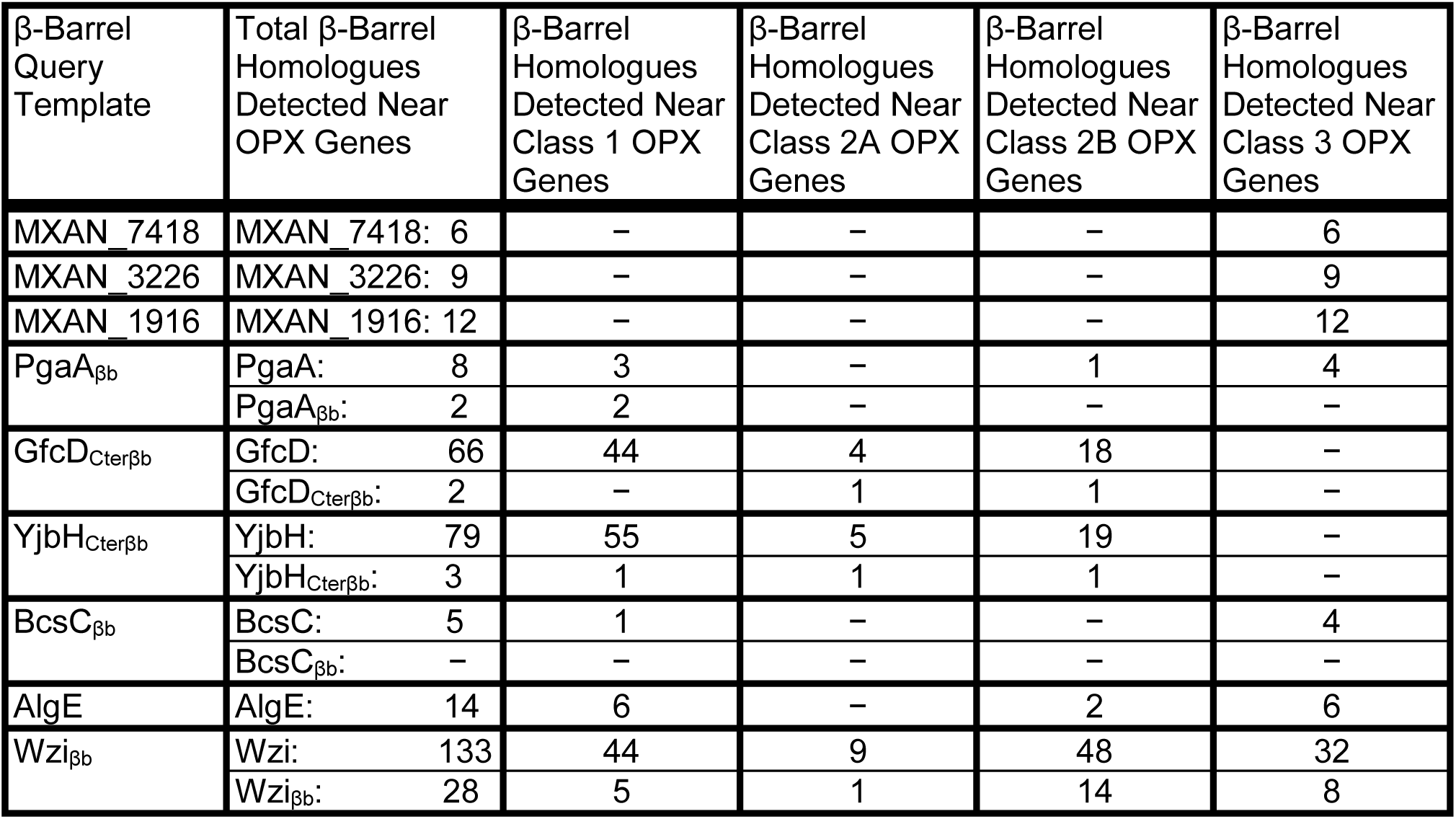
β-barrels identified to be syntenic with OPX genes in the REP dataset.

| β-Barrel Query Template | Total β-Barrel Homologues Detected Near OPX Genes | β-Barrel Homologues Detected Near Class 1 OPX Genes | β-Barrel Homologues Detected Near Class 2A OPX Genes | β-Barrel Homologues Detected Near Class 2B OPX Genes | β-Barrel Homologues Detected Near Class 3 OPX Genes |
| --- | --- | --- | --- | --- | --- |
| MXAN_7418 | MXAN_7418: 6 | – | – | – | 6 |
| MXAN_3226 | MXAN_3226: 9 | – | – | – | 9 |
| MXAN_1916 | MXAN_1916: 12 | – | – | – | 12 |
| PgaA<sub>βb</sub> | PgaA: 8 | 3 | – | 1 | 4 |
|  | PgaA<sub>βb</sub>: 2 | 2 | – | – | – |
| GfcD<sub>Cterβb</sub> | GfcD: 66 | 44 | 4 | 18 | – |
|  | GfcD<sub>Cterβb</sub>: 2 | – | 1 | 1 | – |
| YjbH<sub>Cterβb</sub> | YjbH: 79 | 55 | 5 | 19 | – |
|  | YjbH<sub>Cterβb</sub>: 3 | 1 | 1 | 1 | – |
| BcsC<sub>βb</sub> | BcsC: 5 | 1 | – | – | 4 |
|  | BcsC<sub>βb</sub>: – | – | – | – | – |
| AlgE | AlgE: 14 | 6 | – | 2 | 6 |
| Wzi<sub>βb</sub> | Wzi: 133 | 44 | 9 | 48 | 32 |
|  | Wzi<sub>βb</sub>: 28 | 5 | 1 | 14 | 8 |

Together, these data reveal intriguing architectural similarities between β-barrel porin modules from synthase-dependent polymer export pathways and those implicated in myxobacterial Wzx/Wzy-dependent secretion, as well as analogous or ABC transporter-dependent pathways in diverse bacteria, all previously unreported associations.

### WzpX, WzpS, and WzpB are (respectively) integral OM β-barrel EPS-, MASC-, and BPS-pathway porins

To examine the structural suitability of the WzaX/S/B co-occurring β-barrels for the respective translocation of EPS/MASC/BPS, and since no full-length template structure could be identified, we employed the deep-learning approach provided by AlphaFold2 to generate a tertiary structure model. AlphaFold2 employs information from evolutionarily-coupled amino acids as well as templates with structural homology to fold a polypeptide sequence using an iterative process (Jumper *et al*., 2021). Consistent with the above-described data (**Supplementary Figs. 2,3,4**), MXAN_7418, 3226, and 1916 were all predicted to form 18-stranded β-barrels with sizeable central cavities, with respective barrel heights of 33, 32, and 33 Å (**Fig. 5A**). As molecular dynamics simulations typically calculate the hydrophobic thickness of asymmetric OM bilayers to be ∼40% of their total solvated thickness (Pavlova *et al*., 2016), based on our measured *M. xanthus* OM thickness of 69.8 ± 1.8 Å (**Fig. 4A**), an approximated hydrophobic thickness of ∼28 Å would indeed by traversable by the proposed MXAN_7418, MXAN_3226, and MXAN_1916 tertiary structures.

**Figure 5.**
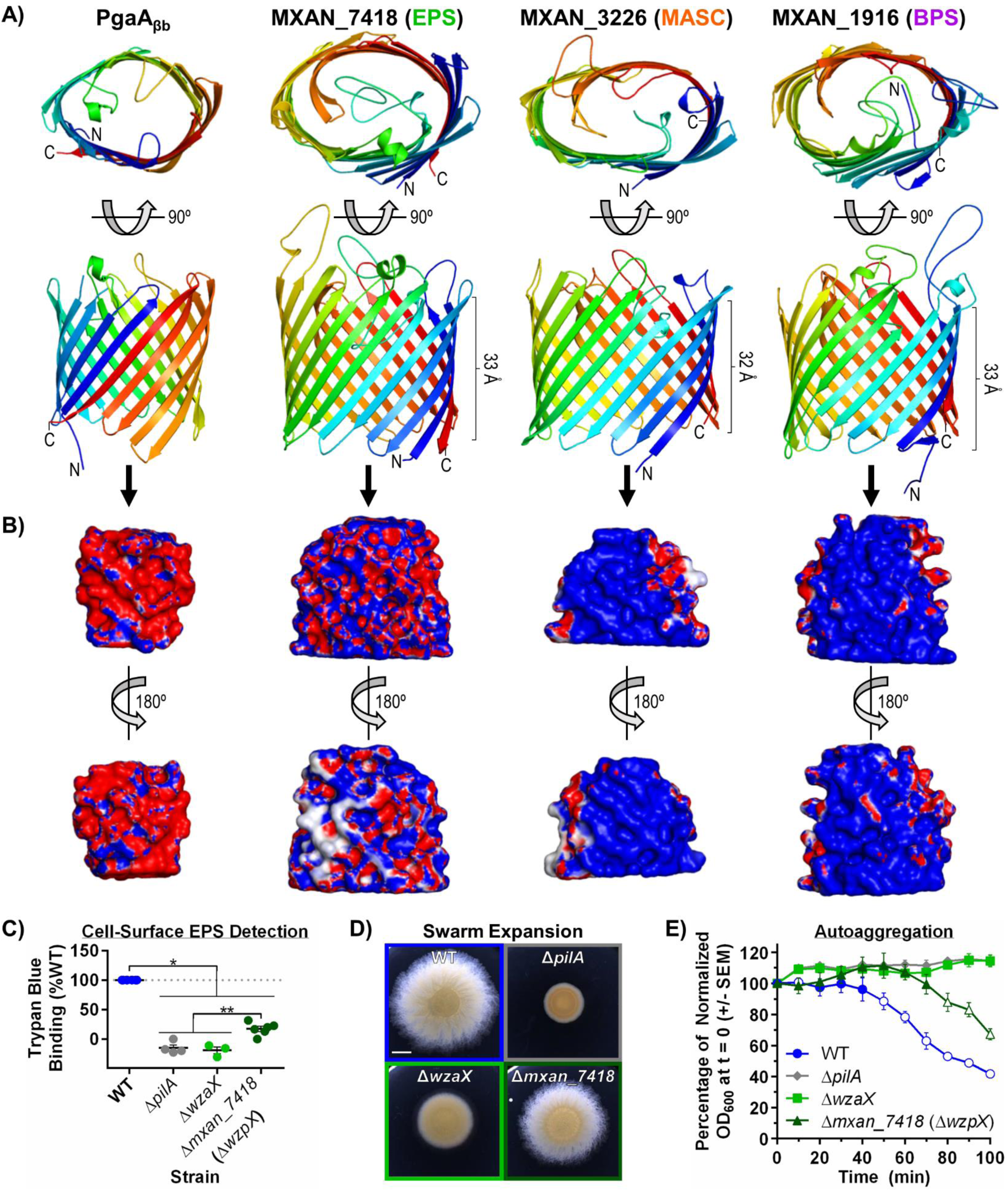
OPX-companion β-barrel structures. **A)** Tertiary structure models (top and front views) for MXAN_7418, MXAN_3226, and MXAN_1916, as generated using deep learning via AlphaFold2, as well as the PgaA C-terminal domain (aa 513–807) X-ray crystal structure (PDB: 4Y25) (Wang *et al*., 2016). Structures are coloured with a spectrum, from the N-terminus (*blue*) to the C-terminus (*red*), and depicted with smooth loops. **B)** Front and back views of the interior spaces of the β- barrels depicted in Panel A overlaid with the electrostatic character of the residues contacting the lumenal volume, as generated via HOLLOW (Ho & Gruswitz, 2008). Surfaces have been colored according to charge, from blue (positive, +5 kT/e) to white (uncharged/hydrophobic), to red (negative, −5 kT/e). **C)** Trypan Blue dye-binding for *M. xanthus* DZ2 WT (n = 6), Δ*pilA* (n = 4), Δ*wzaX* (n = 3), and Δ*mxan_7418* (i.e. Δ*wzpX,* n = 4) to probe cell-surface EPS levels. Mean values are indicated (+/− SEM), with each biological replicate data point indicated. Means of all mutants were significantly lower than WT, while that of Δ*mxan_7418* was significantly higher than either Δ*pilA* or Δ*wzaX*, as calculated via Student’s T-test (*p* < 0.05). **D)** T4P-dependent swarm expansion of strains tested in Panel C. Scale bar: 4 mm. **E)** Auto-aggregation profiles of strains tested in Panel C for cells resuspended in CYE rich medium at an initial OD_600_ of 1.0. Mean values (n = 3) are indicated +/− SEM. Open plot points: no statistically significant difference in mean relative to Δ*wzaX* at a given time point. Closed plot points: statistically significant difference of means relative to Δ*wzaX* at a given time point. Significance was evaluated via Student’s T-test (*p* ≤ 0.05).

We subsequently used HOLLOW to probe the lumenal volume of the EPS/MASC/BPS- cluster β-barrels via filling of the internal space with dummy atoms to generate a cast of the void space, after which the electrostatic potential of the contacting β-barrel surface was overlaid. To validate this approach, we first probed the internal electrostatics of the PgaAβb template structure, revealing a highly anionic interior (**Fig. 5B**), consistent with passage of the cationic PNAG polymer through the lumen of the barrel. BPS was previously discovered to be a randomly-acetylated anionic repeating tetrasaccharide, with the distal three sugars of each repeat constituted by mannosaminuronic acid (ManNAcA) units (Islam *et al*., 2020). Therefore, the cationic charge character of the MXAN_1916 lumen (**Fig. 5B**) is indeed suitable for passage of its associated HMW BPS polymer. While the chemical structures or exact compositions of MASC or EPS are not known, isolated spore coat material was found to contain GalNAc chains with potential glucose (Glc) and glycine decorations (Holkenbrink *et al*., 2014). As the interior of the MXAN_3226 β-barrel is cationic (**Fig. 5B**), this suggests that MASC may have a net-anionic charge character, as contributed via as-yet-unidentified sugars and/or chemical modifications. EPS composition has been probed across four investigations (Islam *et al*., 2020, Behmlander & Dworkin, 1994, Gibiansky *et al*., 2013, Sutherland & Thomson, 1975), with Ara, Gal, GalNAc, Glc, GlcN, GlcNAc, Man, ManNAc, Rha, and Xyl having been identified (depending on the publication); however, none of these sugars are highly charged, which is consistent with the more neutral character of the MXAN_7418 interior (compared to that of either MXAN_1916 or MXAN_3226) (**Fig. 5B**).

To probe the implication of Wzx/Wzy-dependent pathway-associated β-barrels in polymer secretion, we next set out to better understand their physiological contexts. RNAseq analysis previously detected the transcripts encoding MXAN_7418 and MXAN_1916 in vegetative cells, as well as MXAN_3226 in developmental cells, indicating that all three β-barrels are indeed expressed over the course of the *M. xanthus* lifecycle (Muñoz-Dorado *et al*., 2019, Sharma *et al*., 2021).

Furthermore, MXAN_1916 was detected in proteomic screens of biotinylated surface-exposed proteins, and MXAN_1916 and MXAN_3226 were both detected in OMV samples from vegetative cells (Kahnt *et al*., 2010). Importantly, the MXAN_3226 β-barrel was already shown to be an essential part of the MASC pathway as its respective *M. xanthus* deletion-mutant strain was found to be deficient in sporulation and MASC production (Holkenbrink *et al*., 2014). To examine effects of β-barrel deletion on EPS levels in vegetative cells, we first generated a Δ*mxan_7418* chromosomal deletion mutant strain. Since retention of Trypan Blue has become a well-established indicator for the presence of EPS on the surface of *M. xanthus* cells, we next compared the dye-binding capacity of Δ*mxan_7418* cells versus EPS-pathway OPX^−^ (Δ*wzaX*) and T4P^−^ (Δ*pilA*) cells, both known to be defective in EPS production. Relative to WT cells, absence of the EPS-pathway β-barrel resulted in an 83% loss of Trypan Blue retention by Δ*mxan_7418* cells (**Fig. 5C**), indicating a severe reduction in the amount of cell-surface EPS in the β-barrel mutant, consistent with cell-surface EPS deficiencies previously probed in Δ*wzxX*, Δ*wzyX*, Δ*wzcX*, Δ*wzeX*, and Δ*wzaX* EPS-pathway mutants (Islam *et al*., 2020).

Compared to the baseline readings in the Δ*wzaX* EPS-pathway OPX^−^ mutant strain, Δ*mxan_7418* cells displayed marginally higher levels of Trypan Blue binding (**Fig. 5C**). These results are consistent with EPS-pathway β-barrel deficiency principally impacting polymer export to the cell surface, as opposed to both polymer assembly and export being compromised in the absence of EPS-pathway OPX proteins. To probe whether residual quantities of EPS were indeed present on the surface of Δ*mxan_7418* cells, we compared swarm expansion on solid medium as well as auto- aggregation in liquid medium; though both phenotypes are multifactorial, each requires T4P engagement with cell-surface EPS. Relative to WT, Δ*mxan_7418* swarm expansion was reduced, with this phenotype even more pronounced in Δ*wzaX* swarms (**Fig. 5D**). Similarly for auto- aggregation testing in rich medium, consistent with previous findings, WT cells steadily clumped together and sedimented in the cuvette, whereas both Δ*pilA* and Δ*wzaX* cells did not (**Fig. 5E**). Cells of the Δ*mxan_7418* mutant remained in suspension analogous to Δ*wzaX* cells for ∼75% of the assay, after which they began to slowly sediment (**Fig. 5E**), suggesting that cell-surface EPS had eventually accumulated to a sufficient threshold to support T4P-mediated clumping in liquid. Taken together, these data indicate that while minimal amounts of EPS can reach the cell surface through as-yet- undetermined means (see Discussion for further comment), MXAN_7418 serves as the principal trans-OM conduit for EPS export in *M. xanthus*.

Ultimately, the findings detailed in this investigation support a model for polysaccharide export in Wzx/Wzy-dependent pathways lacking an integral OM OPX protein — as represented by the independent EPS, BPS and MASC pathways in *M. xanthus* — in which the final polymer must pass through an integral OM β-barrel porin for efficient secretion to the cell exterior. For these reasons, as well as the long-established naming convention for Wzx/Wzy-dependent pathways in bacteria (Reeves *et al*., 1996), we propose the designation Wzp (i.e. <u>Wz p</u>orin) for the newly- identified component of these secretion systems (**Fig. 6**).

**Figure 6.**
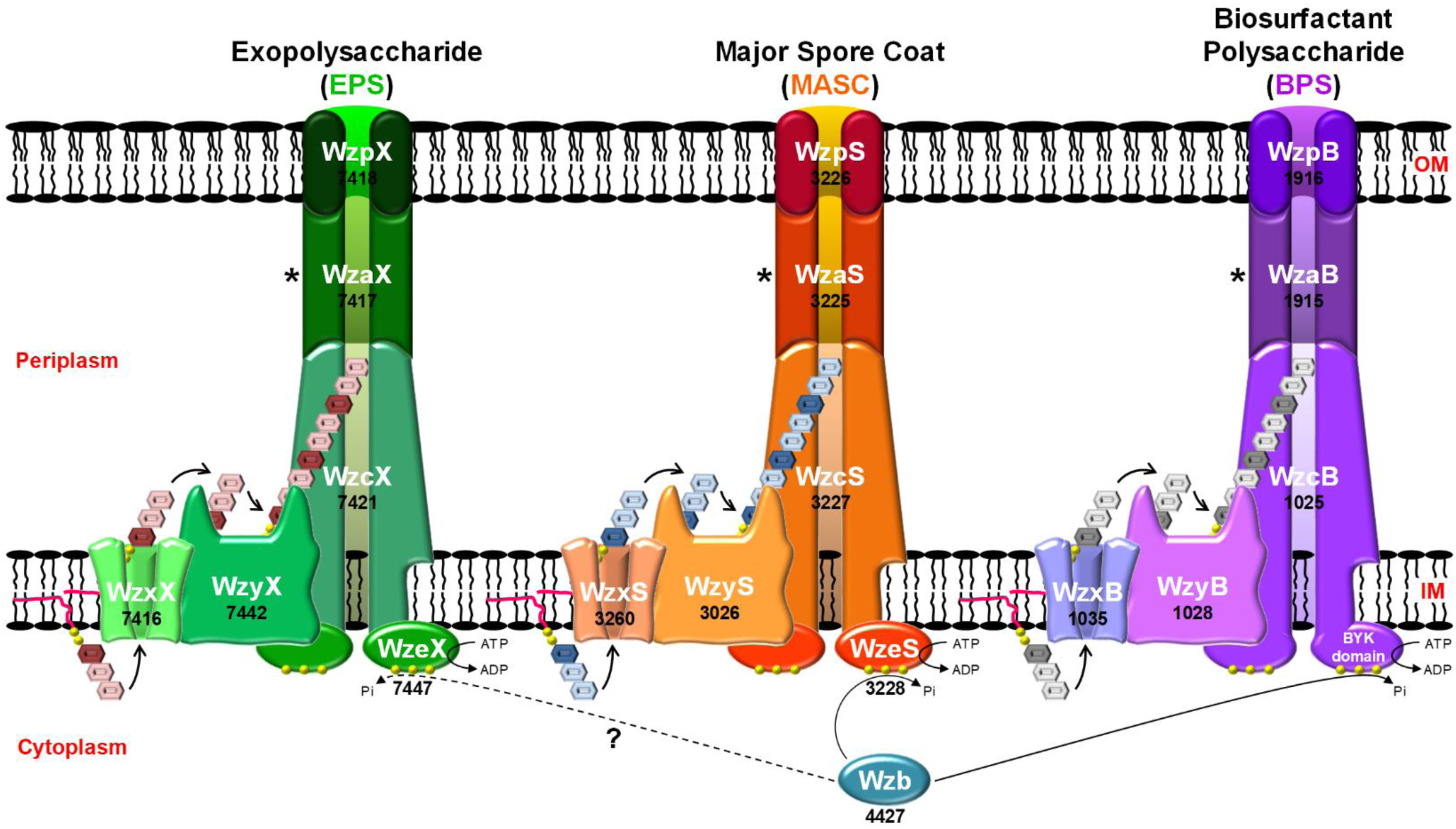
Wzx/Wzy-dependent polysaccharide assembly-and-secretion pathways in *M. xanthus*. In these schematics, the WzaX/S/B proteins have been depicted in a linking capacity between the apex of the WzcX/S/B PCP periplasmic domains and the periplasmic opening of the integral OM WzpX/S/B β-barrel porins identified herein. However, (i) the exposure of the EPS/MASC/BPS polymers to the periplasmic space as each transits between the IM and OM components of each system, and (ii) the exact role(s) of the WzaX/S/B proteins in *M. xanthus* polymer translocation (Supplementary Fig. S6), remain open questions for each pathway. To denote these uncertainties, this stage of the transport cycle has been marked with an asterisk (*).

## DISCUSSION

Knowledge of the terminal component through which secreted polysaccharides exit a bacterial cell is crucial for the development of targeted antimicrobial agents that could be used to inhibit this process (Kong *et al*., 2013). From the data presented herein, we have provided evidence that bacterial OPX proteins fall into one of three distinct structural classes, the first two of which have either the demonstrated (Class 1) or predicted (Class 2) capacity to span the OM, whereas the third (Class 3) is missing any such domains. Instead, as demonstrated by the Class 3 OPX WzaX/S/B proteins from *M. xanthus*, these Wzx/Wzy-dependent pathway proteins are genomically and functionally paired with a complementary integral OM-spanning β-barrel porin similar to that required for PNAG export in synthase-dependent pathways.

The Class 1 OPX protein Wza*Ec* is the most extensively-characterized OPX protein with respect to structure–function relationships. The orientation of Wza*Ec* in the OM was established through introduction of a FLAG tag on the Wza*Ec* C-terminus, allowing for a FLAG epitope to be detected on the cell surface with α-FLAG antibodies (Dong *et al*., 2006). Similar introduction of a C- terminal His6 epitope tag resulted in a partially functional Wza*Ec*-His6 construct that was able to restore K30 Group 1 CPS biosynthesis to ∼20% of the level restored by an untagged Wza*Ec* construct following expression *in trans* (Nesper *et al*., 2003). Various C-terminal α-helix truncations also abolished Wza*Ec* function (Ford *et al*., 2009). This indicates that the OM-spanning domain of Wza*Ec* can be functionally-sensitive to structural perturbation. In lipid bilayers, purified Wza*Ec*-His6 was able to form 2D octameric ring-like crystal arrays (Beis *et al*., 2004), suggestive of channel formation, with this arrangement confirmed by the 3D X-ray crystal structure (Dong *et al*., 2006).

Through introduction of the photo-crosslinkable synthetic amino acid *p*-benzoyl-L-phenylalanine (*p*Bpa) at various sites in Wza*Ec*, Nickerson and colleagues elegantly demonstrated that K30 CPS polymers could be trapped in the lumen of the translocon (Nickerson *et al*., 2014), confirming that polysaccharides do indeed pass through Wza*Ec* during secretion. Specifically, substitutions at certain sites within periplasmic D2 (**Fig. 1A**) were able to maintain translocon functionality as well as form crosslinks with polymers. Site-specific *p*Bpa substitutions within periplasmic D1 rendered Wza*Ec* non-functional. Conversely, *p*Bpa-substituted positions in periplasmic D3, and more importantly at the kink in the OM-traversing α-helical D4, maintained Wza*Ec* functionality, but were unable to form detectable intermolecular crosslinks upon UV exposure (Nickerson *et al*., 2014). As such, the transit of secreted polymer through the α-helical D4 Wza_C domain pore could not be demonstrated via this technique. However, purified Wza*Ec* was shown to stably insert into planar lipid bilayers and form electro-conductive channels, with site-specific amino acids substitutions confirming that ion flow does indeed proceed via the D4 pore formed by the OM-spanning α-helix (Kong *et al*., 2013).

Synthetic peptides corresponding to the native Wza*Ec* D4 α-helix sequence were also able to spontaneously insert into such bilayers, but formed unstable pores; however, modification of the native D4 sequence through consensus generation (based on 94 closely-related sequences) yielded an optimized peptide (**Fig. 2B**) that could spontaneously insert into bilayers and form stable pores (Mahendran *et al*., 2017). Given the primary structure diversity amongst OM-spanning α-helices uncovered herein for both Class 1 and Class 2B OPX proteins (**Fig. 2B,C**), additional optimization of synthetic peptide sequences should be possible to further improve spontaneous membrane insertion, self-assembly, and α-barrel pore stability.

The Class 2B OPX protein KpsD*Ec* has long been known to be essential for Group 2 CPS export. When expressed in isolation, KpsD*Ec* was shown to reside in the periplasm (Silver *et al*., 1987). However, upon expression of KpsD*Ec* along with its cognate IM-localized PCP KpsE (which extends into the periplasm), KpsD*Ec* was also detected in IM and OM fractions of lysed cells (Arrecubieta *et al*., 2001). Intriguingly, OM-localized KpsD*Ec* is detected as a multimer, whereas the minimally-detected IM-localized KpsD*Ec* is present as a monomer, consistent with the adoption of quaternary structure by KpsD*Ec* at the site of polysaccharide egress from the cell (Sande *et al*., 2019). Furthermore, immunolabelling of intact *E. coli* cells using α-KpsD*Ec* antiserum resulted in the detection of KpsD*Ec* epitopes at the cell surface (McNulty *et al*., 2006), suggesting that a portion of KpsD*Ec* was indeed surface-accessible. The detection herein of structural homology of the extreme KpsD*Ec* C-terminus to the OM-spanning domain of Wza*Ec* further supports the contention that a part of KpsD*Ec* is able to interact with, and span, the OM bilayer, albeit in a conditional manner. The 24- residue length of the KpsD*Ec* C-terminal α-helix is well within the threshold of 20 amino acids required to span the hydrophobic core of a membrane bilayer for α-helical integral membrane domains (Baeza-Delgado *et al*., 2013). Analogous to Wza*Ec*, truncation of the KpsD*Ec* C-terminus by 11 amino acids abrogated protein function (Wunder *et al*., 1994). A potential OM-spanning α-helix is indeed a conserved property of the Class 2B OPX proteins identified in this study. Despite the detectability of KpsD*Ec* at the cell surface with α-KpsD*Ec* antibodies, α-His-tag antibodies were unable to label the surface of cells expressing a KpsD*Ec* variant encoding a C-terminal His6 affinity tag (McNulty *et al*., 2006); in this instance, the highly-cationic nature of the affinity tag may have impeded its translocation across the hydrophobic OM bilayer, thus inhibiting immunodetection.

Given that KpsD*Ec* by itself does not intrinsically associate with the OM (Silver *et al*., 1987), this may suggest that the protein can become directly inserted into the OM (as opposed to requiring OM insertion via the Bam/Tam or Lol machinery). In this manner, KpsD*Ec* could indeed function as the terminal piece of the Group 2 capsule secretion machinery.

Historically, Group 2 CPS secretion across the OM had been suggested to implicate integral OM β-barrel porins (Whitfield & Valvano, 1993, Bliss & Silver, 1996), however such a model has fallen out of favour given that no such β-barrels have ever been detected in or near related Kps synthesis clusters. This would be consistent with an ability of KpsD*Ec*-like Class 2B OPX proteins to traverse the OM, thus not requiring any downstream piece of export machinery in certain organisms. However, this scenario may not be absolute, as numerous instances of β-barrel proteins encoded near both Class 2A and Class 2B (as well as Class 1 and Class 3) OPX genes were uncovered herein, suggesting that integral OM porins may play an important role in non-synthase-dependent secretion in diverse organisms. Though not all Class 2 OPX proteins identified in our investigation were matched with a nearby β-barrel, our synteny analysis window was limited to +/− 10 genes from each OPX gene and would thus not have captured candidate porins elsewhere in the genome. As a case- in-point, the *M. xanthus* EPS, MASC, and BPS clusters contain respective insertions of >18 kbp, >223 kbp, and > 1 Mbp that separate constituent members of the same cluster, which in the latter results in extraordinary genomic distance between *wzaB*-*wzpB* and the remainder of the BPS assembly genes (Islam *et al*., 2020). Moreover, as our synteny analyses were limited to 9 query sequences (8 with similarity to PgaAβb), this does not preclude the presence of other/more distantly-related β-barrels near “unmatched” OPX proteins. However, in the absence of specific templates with which to search, such an analysis was beyond the scope of the current investigation.

The identification of “OPX” proteins (an obvious misnomer) in Gram-positive bacteria, but specifically those of Class 3 architecture, point to a broadly-conserved periplasmic function for these proteins, likely through interfacing with the periplasmic domains of PCP proteins in most organisms. However, as dedicated peptidoglycan-spanning polysaccharide export channels have yet to be identified in Gram-positive bacteria, any role for Class 3 OPX proteins in these systems with regards to interactions with a secretion pore of some sort would be unfounded speculation. In *M. xanthus* cells, WzaX/S/B Class 3 OPX protein deficiency does not lead to visible accumulations of polymeric material in the periplasm (Saïdi *et al*., 2021) suggesting that EPS/MASC/BPS polymer assembly via Wzx/Wzy-dependent pathways does not indiscriminately continue in these mutant backgrounds; this is a similar observation to that for Wza*Ec* Class 1 OPX deficiency in *E. coli* cells (Nesper *et al*., 2003). Such material from ABC transporter-dependent synthesis does however accumulate in the periplasm of KpsD*Ec*-deficient Class 2B OPX-mutant cells (Wunder *et al*., 1994, Bliss & Silver, 1996).

For myxobacterial Class 3 OPX proteins, a clear partnership has now been demonstrated between these secretion-pathway components and integral OM β-barrel porins illustrating the requirement of the latter for HMW polysaccharide export across the OM. The WzpS (MXAN_3226) β-barrel was already shown to be essential for MASC secretion, but its function in the MASC transport cycle was not known at the time (Holkenbrink *et al*., 2014). In the current investigation, we have shown that the properties of WzpB (MXAN_1916) make it suitable for BPS translocation across the OM. Furthermore, we have herein demonstrated that WzpX (MXAN_7418) functions as the principal export conduit for EPS across the OM to the cell surface. Intriguingly, EPS still appears to be assembled to a certain degree in mutant cells lacking WzpX (given the residual amount detected on the cell surface), as opposed to cells lacking WzaX which manifest an even more robust EPS^−^ phenotype. This may indicate that while lack of the OPX component in the EPS pathway results in a severe reduction (or shutdown) of EPS production, in cells lacking WzpX, other β-barrel porins (e.g. WzpS and/or WzpB) may be able to inefficiently cross-complement the deficiency in the EPS-pathway machinery, resulting in residual EPS localization to the cell surface. Part of this inefficiency could arise from the cationic natures of the WzpS and WzpB barrel interiors not serving as suitable conduits for a more neutral EPS polymer.

The presence of a Wza_C domain in a Class 1 or 2B OPX protein is not mutually exclusive to the presence of an integral OM β-barrel protein encoded nearby, considering that such barrels were found to be encoded near genes representing all three OPX protein classes across a range of bacterial genomes. The GfcD protein is particularly intriguing given the GfcE protein also encoded by the Group 4 CPS export locus (Peleg *et al*., 2005). While GfcD contains a PgaAβb-like polysaccharide secretion component with an attached SLBB-like domain (as part of the GfcD C-terminal module) (**Supplementary Fig. 5B**), GfcE is a functional Class 1 OPX paralogue of Wza*Ec*; this was evidenced by the ability of GfcE (formerly YccZ/Wza22min) expressed *in trans* to partially restore *E. coli* K30 CPS production in a mutant lacking Wza*Ec* (Drummelsmith & Whitfield, 2000). GfcE possesses a complete Wza_C domain, and as such would be expected to span the OM in a Wza*Ec*-like manner. One possibility could be that the Wza_C domain of the GfcE OPX protein may be required to properly interact/organize around the GfcDCterβb polysaccharide secretion module. Furthermore, the presence of a putative FapF amyloid secretion β-barrel fused to the same polypeptide as that of a PgaAβb-like module typically associated with polysaccharide secretion raises an interesting possibility. Amyloid proteins are frequently secreted by bacteria in order to stabilize biofilm matrices composed largely of secreted polysaccharides (Erskine *et al*., 2018). Thus, in Group 4 capsules, secretion of amyloidogenic polypeptides (via the GfcD FapF-like N-terminal module) could help to stabilize the polysaccharide component of the capsule structure and/or anchor the CPS to the cell surface.

### Ideas and Speculation

For Class 3 OPX proteins, the lack of OM-spanning domains and the (relatively) small size of these proteins (compared to other OPX classes), present a dilemma regarding the mechanism of transit across the periplasm for polymers produced by these systems (**Supplementary Fig. 6**). As a case-in-point, the thickness of the *M. xanthus* periplasm was measured to be 327 Å (**Fig. 4A,B**), while the periplasmic domain of the BPS-pathway WzcB PCP-2B protein was estimated to extend up to ∼165 Å into the periplasmic space. Coupled with the maximum possible height of ∼62 Å for one unit of OPX protein WzaB, together this only accounts for ∼227 Å of periplasmic thickness, leaving ∼100 Å of periplasmic height unaccounted for (**Supplementary Fig. 6**), compared to standard models of polysaccharide secretion (**Fig. 1A**). Furthermore, high-confidence co-evolving amino acids between WzcB and WzaB localize to the apex of the PCP and the base of the Poly_export domain of the Class 3 OPX protein (respectively) (**Supplementary Fig. 6, Supplementary Table S5**), heavily implying the presence of a conserved interaction interface between the two proteins. With the assumption of pathway specificity for each *M. xanthus*Class 3 OPX protein — as evidenced by lack of cross-complementation in single-OPX-knockout strains (Islam *et al*., 2020) — and using components of the BPS pathway as examples (**Fig. 6**), several potential models for trans-envelope transit can thus be proposed:

(i) Model 1: As the polymer exits the WzcB cavity in the periplasm (following WzyB- mediated polymerization), it passes through a periplasmic WzcB-associated single-layer WzaB oligomer. Once past this point, the polymer would have to independently locate its cognate integral OM WzpB β-barrel porin in order to reach the outside of the cell. This presumes the polymer is exposed to the periplasm for a substantial portion of its trip between the IM and OM. In such a model, polymer export might still be possible in the absence of the cognate OPX (as long as this absence does not impact polymer assembly). However, since this is not the case in *M. xanthus*, it would argue against this model.
(ii) Model 2: This is a variation of Model 1 in which single-layer periplasmic oligomers of WzaB are located both on the apical point of the PCP octamer as well as the proximal face of the integral OM WzpB β-barrel porin. In this manner, *M. xanthus* OPX-mediated substrate specificity would exist at both ends of the transport process, but again the polymer could be exposed to the periplasm during stages of the transport cycle bridging the OPX proteins. Nonetheless, this model provides a solution to the question of “targeting” of the nascent polymer to the proper OM-spanning apparatus. However, this model also presumes a constant presence of Class 3 OPX proteins associated with the OM, as well as specific interactions between the periplasmic OPX WzaB and the integral OM porin WzpB. At the moment, this is not bolstered by high-confidence evolutionary couplings data between the two proteins (**Supplementary Table S5**), but this may be partially due to an insufficient number of barrel homologues with which to fully probe coevolution. However, as OPX proteins are not detected in surface-biotinylated or OMV samples from *M. xanthus*, supporting evidence for this concept is lacking.
(iii) Model 3: The periplasmic distance of 327 Å could be approximately accounted for by ∼165 Å of WzcB PCP-2B periplasmic domain height, followed by OPX oligomers of WzaB stacked in duplicate (∼62 Å × 2 = ∼124 Å) or triplicate (∼62 Å × 3 = ∼186 Å), depending on the packing arrangement of the oligomers. Along with the PCP channel, such architecture could be envisaged to form a protected channel lumen spanning from the IM to the OM, precluding exposure of the polymer to the periplasm. This would also abrogate any “targeting” issues of the polymer to the WzpB β-barrel secretin. In this case, HMW oligomers of *M. xanthus* OPX proteins would be expected in the periplasm, which could possibly co-precipitate with IM and/or OM fractions. Once again, data to support this contention awaits further experimentation.
(iv) Model 4: As the nascent polymer emerges from the PCP-2B opening, the polymer interacts with copies of WzaB at the apex of the PCP-2B periplasmic domain, allowing WzaB to bind the polymer and detach from the PCP. As additional polymer elongation occurs, more units of WzaB are able to bind further down the polymer. In this manner, WzaB bound to the polymer would serve as a type of targeting chaperone which preferentially directs the periplasmic polymer to its cognate WzpB β-barrel porin in the OM. Once a given WzaB has reached the porin, it would disengage from the polymer and be able to undergo subsequent rounds of binding in the periplasm to the nascent polymer. Such a mechanism would afford a certain degree of protection to the translocating polymer against the periplasmic environment, as well as provide a mechanism for targeting of the polymer to the specific β-barrel machinery needed for transport across the OM. This would also explain why the Class 3 OPX proteins WzaX, WzaS, and WzaB were not detected in screens of surface-biotinylated proteins or those identified in OMV fractions (from vegetative and developmental cells), i.e. that OM association of WzaX/S/B (via interaction with WzpX/WzpS/WzpB) is of a more transient nature.

Exposure of a sugar polymer to the periplasm as it transits between the IM and OM is a common occurrence in synthase-dependent systems such as those involved in alginate, cellulose, and PNAG assembly-and-export (Whitney & Howell, 2013). Recently, Group 2 CPS was also suggested to be exposed to the periplasm at some point during its secretion across the cell envelope (Liston *et al*., 2018). In these systems, periplasmic enzymes are able to access the transiting polymer and introduce various modifications including sugar epimerization and/or (de)acetylation. Such a mechanism may be in place for the *M. xanthus* BPS polymer, for which the secreted form displays random acetylation (Islam *et al*., 2020). In synthase-dependent pathways, TPR domains (either standalone or attached) extend into the periplasm from the β-barrel porin and are proposed to interact with these chain-modifying enzymes (Whitney & Howell, 2013). Of note, numerous OPX proteins were found to be encoded near β-barrel genes for full-length PgaA and BcsC homologues complete with the respective TPR architectures (**Table 1**). In these systems, questions arise as to the functional relationships between periplasmic OPX proteins and integral OM β-barrel porins with TPR domains potentially occluding access of the OPX protein to the periplasmic face of the porin.

Ultimately, this study reveals OPX-protein complexities in diverse organisms that differ from the well-studied Wza*Ec* protein, which will further our understanding of the mechanism of sugar polymer export in bacterial cells. Moreover, updated genomic, structural, and functional knowledge of the terminal step in the polysaccharide secretion pathway will enable researchers to selectively develop novel antimicrobial compounds targeted to blocking bacterial polymer secretion from the outside, thus bypassing any requirements for access to the cell interior to compromise the viability of a bacterial cell.

## MATERIALS AND METHODS

### Protein structure analysis & modelling

Given the high fold-recognition equivalence between WzaX/S/B and Wza*Ec*, the tertiary structure of each *M. xanthus* OPX protein was modelled against the Wza*Ec* template (PDB: 2J58) using MODELLER. For β-barrel proteins MXAN_7418/3226/1918 that lacked a suitable full-length structural template, protein structure models were computed using deep learning and artificial intelligence via AlphaFold2 (Jumper *et al*., 2021). Multiple sequence alignment entries were generated using 606, 251 and 101 unique sequences, respectively for MXAN_1916, MXAN_3226 and MXAN_7418, with the program run for 5 independent prediction models, leading to convergence after 3 recycling iterations. For proteins GfcD and YjbH, deposited AlphaFold2-generated structures were mined from the UniProt entries for P75882 and P32689, respectively. HOLLOW (Ho & Gruswitz, 2008) was used to generate internal volume casts of the MXAN_7418/3226/1918 β- barrels, followed by overlaying of the solvent-accessible electrostatic potential contributed by amino acids in contact with the internal volume, as calculated using PDB2PQR and APBS (Propka pH 7.0, Swanson force field, ±5 kT/e). All protein structures were visualized and rendered in PyMol.

Evolutionarily-coupled amino acids within the same protein were analyzed using RaptorX-Contact (Wang *et al*., 2017), while evolutionarily-coupled residues between two proteins were determined using RaptorX-ComplexContact (Zeng *et al*., 2018) (http://raptorx.uchicago.edu/). Protein contact maps were displayed in GraphPad.

### Identification of OPX proteins

Three types of datasets i.e., 61 order Myxococcales genomes (MYXO), 3662 reference and representative bacterial genomes (REP; downloaded on Dec 7, 2021), and non-redundant NCBI database (NR) (371 327 556 proteins at 100% identify as of June 10, 2021) were downloaded from NCBI. Pfam domains attributed to PF02563 [Poly_export], PF10531 [SLBB], PF18412 [Wza_C], and PF06251 [Caps_synth_GfcC; used here as GfcC] were extracted from the Pfam-A v34.0 database (Mistry *et al*., 2021) (downloaded: March 24, 2021) and a reduced combined profile database was created. These functional domains were identified by scanning all three types of datasets (MYXO, REP, and NR) using offline hmmscan (Potter *et al*., 2018) against the created database with an E-value cutoff of 1×10^−5^. The resultant files were parsed using hmmscan-parser.sh, sorted and arranged in the form of protein architecture using in-house scripts. Based on the identified domains per protein per dataset, three primary clusters were curated. All identified proteins within these primary clusters were subjected to fold-recognition analysis using HHpred (Söding *et al*., 2005) against the database of two proteins (PDB: 2J58 [i.e. Wza*Ec*] and PDB: 3P42 [i.e. GfcC]; extracted from PDBmmCIF70 downloaded on Nov 19, 2021) using “-p 5 -Z 500 -loc -z 1 -b 1 -B 500 -all -id 35 -ssm 2 -sc 1 -seq 1 -dbstrlen 10000 -norealign -maxres 32000” parameters. HHpred raw data was parsed using in-house scripts to generate the architecture of each protein in terms of non-overlapping homologous regions to 2J58 and 3P42. To identify the type of signal peptide and the cleavage site location, each OPX protein was subject to SignalP 6.0 analysis (Teufel *et al*., 2022) and to predict the membrane topology of the proteins, TMHMM (Server v. 2.0) (Krogh *et al*., 2001) was used.

Based on HHpred analysis, we also investigated the secondary structure-based homology of primary cluster proteins with a Wza_C segment (aa 326 – 359; 34 amino acid length) in PDB 2J58. If a protein was showing secondary structural homology with at least 10/34 amino acids of the Wza_C segment (aa 326 – 359) in 2J58, we considered it as a true Wza_C segment in the respective protein. Proteins with both 2J58- and 3P42-homologous non-overlapping regions were classified as Class 2. Proteins having only 2J58-homologous regions along with a Wza_C segment were classified as Class 1. Other proteins with only 2J58-homologous regions and no Wza_C segment were classified as Class 3. We also generated sequence logos using WebLogo (Crooks *et al*., 2004) (v.2.8) for 2° structure-based homologues of Wza_C segments as identified in Class 1 and Class 2B OPX proteins.

### Synteny analysis of β-barrel query proteins with identified OPX proteins

Along with the identification of the above-mentioned domains and classification of all OPX proteins into three classes, we identified the β-barrel homologues encoded in the vicinity (± 10 genes) of our OPX genes in genomes from both the MYXO and REP databases. We analyzed the various predicted proteomes of both datasets with BLASTp and hmmscan using protein sequences from *M. xanthus* DZ2 (MXAN_7418 [aa 24 – 415], MXAN_3226 [aa 23 – 381], and MXAN_1916 [aa 26 – 421]), *E. coli*K12 (PgaA [aa 511 – 807], GfcD [aa 425 – 698], YjbH [aa 423 – 698], BcsC [aa 785 – 1157], and Wzi [aa 92 – 479], as well as *P. aeruginosa* PAO1 AlgE [aa 33 – 490]. Detected homologues for each query profile were aligned against the truncated and full-length sequences for each query protein using Clustal Omega to probe for the lone presence of a putative β-barrel polysaccharide secretion porin domain versus its presence as part of a multi-domain polypeptide comparable to the native sequence. Hits resembling truncated versions of various β-barrel queries were individually profiled via fold-recognition using the online HHpred bioinformatics toolkit (Zimmermann *et al*., 2018) (https://toolkit.tuebingen.mpg.de/tools/hhpred). Depicted sequence alignments were displayed in GeneDoc with residues colored according to conservation score (out of 10) as indicated in JalView (Waterhouse *et al*., 2009).

### Phylogenetic analysis of OPX proteins

The PF02563 [Poly_export] domain of OPX proteins was used as a phylogenetic marker. The locations of all identified ‘Poly_export’ domains were first extracted, after which extracted sequences were aligned using MUSCLE (with 10 iterations). The resultant alignment was analyzed via FastTree 2.1.10 to generate a maximum likelihood tree of OPX proteins. Tree visualization as well as mapping of the OPX protein classification and taxonomy of each branch were performed via the iTOL web server Version 6.4.3 (Letunic & Bork, 2021).

### Bacterial membrane modeling and intermembrane distance measurements

Desired prokaryotic tomograms were downloaded from the Caltech Electron Tomography Database (https://etdb.caltech.edu/). Forty species were analyzed, each via three separate tomograms. Tomogram inspection and modeling were performed using the IMOD software package (Kremer *et al*., 1996). Tomograms were first oriented in 3D using the IMOD “Slicer” window to identify the central slice through each bacterium. To enhance contrast, 5 layers of voxels were averaged around the section of interest. Model points were then placed along corresponding regions of the OM and IM for a total distance of ∼100 nm. A custom Python script was subsequently used to calculate the intermembrane distance every 0.1 nm along the modeled stretch of membranes. GraphPad was used to prepare plots and carry out correlation analyses.

### Bacterial cell culture

Information on wild-type *M. xanthus* DZ2 (Campos & Zusman, 1975) and isogenic mutant strains analyzed herein can be found in **Table 2**. Strains were grown and maintained at 32 °C on Casitone-yeast extract (CYE) (1% casitone, 0.5% yeast extract, 10 mM MOPS [pH 7.5], 4 mM MgSO4) 1.5% agar (BD Difco) plates or in CYE liquid medium at 32 °C on a rotary shaker at 220 rpm.

**TABLE 2.** *Myxococcus xanthus* strains used in this study.

| Strain | Genotype/Description | Source or Reference |
| --- | --- | --- |
| SI1 | DZ2 (wild type) | (Campos & Zusman, 1975) |
| TM389 | $\Delta pilA$ ($\Delta mxan\_5783$) | (Ducret *et al.*, 2012) |
| TM469 | $\Delta wzaX$ (i.e. $\Delta mxan\_7417/epsY$) | (Ducret *et al.*, 2012) |
| SI93 | $\Delta wzpX$ (i.e. $\Delta mxan\_7418/epsX$) | This study |

### Mutant construction

As previously described (Islam *et al*., 2020), to generate a *M. xanthus* deletion-mutant strain, 500 bp upstream and 500 bp downstream of the target gene were amplified and fused via PCR, subjected to double digestion with restriction enzymes, then ligated into the pBJ114 plasmid. Products were used to transform chemically-competent *E. coli* DH10B via heatshock, after which cells were plated on LB agar supplemented with kanamycin (50 µg/mL) to select for drug-resistant colonies. Successful clones were verified via sequencing. Resultant plasmids were then introduced into *M. xanthus* DZ2 via electroporation. Mutants resulting from homologous recombination of deletion alleles were obtained by selection on CYE agar plates containing kanamycin (100 µg/mL). A second selection was then made on CYE agar plates containing galactose (2.5%) to obtain the final deletion strain, with mutants verified via PCR amplification using flanking primers.

### Trypan blue dye retention

Retention of the dye Trypan Blue was carried out as previously described (Islam *et al*., 2020).

In brief, cells from overnight CYE cultures were first resuspended to OD600 1.0 in TPM buffer. Resuspended cells (or a cell-free blank) (900 µL) were then mixed with Trypan Blue solution (100 µL, 100 µg/mL stock concentration) in a microfuge tube and briefly pulsed (1 s) via vortex mixer. Samples were incubated at room temperature, in a tube rack covered with aluminum foil, atop a rocker platform (1 h) to facilitate dye binding by the cells. Samples were subsequently sedimented (16 000 × *g*, 5 min), after which the top 900 µL of blank or clarified supernatant was transferred to a disposable spectrophotometer cuvette. The cell-free “TPM + Trypan Blue” sample was used to blank the spectrophotometer at 585 nm. The absorbance at 585 nm (A585) was then determined for each clarified supernatant. Finally, absorbance values were normalized to the respective A585 value for the WT of each biological replicate. Sub-zero final values are due to trace amounts of cell debris detected at 585 nm in individual samples in which absolutely no binding of Trypan Blue occurred.

### Phenotypic analyses

Cells from exponentially-growing cultures were harvested and resuspended in TPM buffer (10 mM Tris-HCl, pH 7.6, 8 mM MgSO4 and 1 mM KH2PO4) at a final concentration of OD600 5.0. To study T4P-dependent swarm expansion, this cell suspension (5 μL) was spotted onto CYE 0.5% agar. Plates were incubated at 32 °C (72 h), then imaged with an Olympus SZX16 stereoscope with UC90 4K camera. For T4P-dependent motility, swarm images were captured using the 0.5× objective at 1× zoom, using linear color and darkfield illumination.

### Auto-aggregation testing

The protocol followed has been previously detailed (Saïdi *et al*., 2021). In brief, the turbidity (OD600) of *M. xanthus* CYE cultures (12.5 mL) grown overnight was determined via spectrophotometer. Specific culture volumes were aspirated, then sedimented via microfuge (4000 × *g*, 5 min) so pellet resuspension in 1 mL CYE broth would yield a final OD600 of either 1.0. Broth- resuspended cells were then transferred to a polystyrene spectrophotometer cuvette. Resuspensions were strongly aspirated and ejected in the cuvette (10 s) via p200 micropipette, then immediate read for OD600 (*t* = 0) in a spectrophotometer. Time-course readings of OD600 were taken every 10 min up to 100 min of monitoring. In between readings, cuvettes were left covered and undisturbed on the benchtop in a cuvette box. All OD600 readings were normalized to the OD600 determined at *t* = 0 for each sample.

## ACKNOWLEDGEMENTS

The authors would like to thank Yossef Lopez de Los Santos for protein modelling feedback, Omaima Rebay for cloning assistance, and Joseph Lam for critical reading of the manuscript. A Discovery operating grant (RGPIN-2016-06637) from the Natural Sciences and Engineering Research Council of Canada (NSERC) supported this work in the lab of S.T.I. as well as studentships for F.S. and N.Y.J. F.S., N.Y.J., and R.B. are recipients of graduate studentships from the PROTEO research network. This work was also supported by (i) a DST-INSPIRE Faculty award to G.S. from the Department of Science and Technology (DST), India, a (ii) DST-INSPIRE Fellowship to U.M. from the DST, India, (iii) partial support from the Department of Electronics, IT, BT, and S&T of the Government of Karnataka, India to U.M., A.P., and G.S., and (iv) a David and Lucile Packard Fellowship for Science and Engineering (2019-69645) to Y.-W.C. The funders had no role in study design, data collection and interpretation, or the decision to submit the work for publication. The authors declare that the research was conducted in the absence of any commercial or financial relationships that could be construed as a potential conflict of interest.

## AUTHOR CONTRIBUTIONS

STI and GS conceived of and planned the study.

UM, AP, and GS performed comparative genomics studies. FS and RB generated mutant strains.

FS performed phenotypic, dye-binding, and auto-aggregation analyses. FS and NYJ performed HOLLOW and electrostatics analyses.

STI and AM carried out protein modelling.

MM and GJ designed the periplasmic analysis workflow, with analysis by FS. STI, GS, and FS wrote the manuscript.

STI and GS generated figures.

STI, GS, CC, and YWC contributed personnel and funding support.

---

**Supplementary Figure S1.**
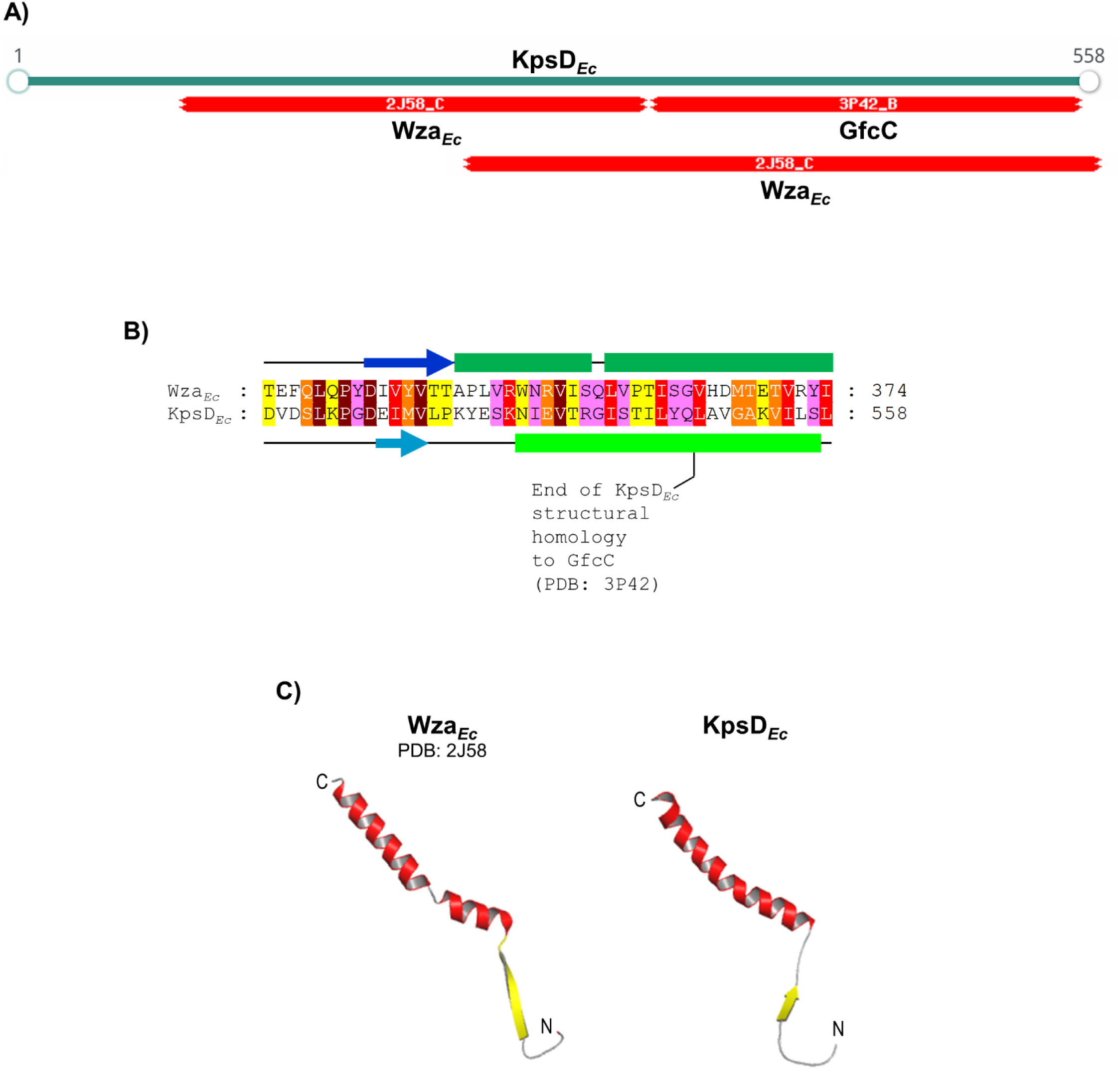
Structural homology between Wza*_Ec_* and KpsD*_Ec_*. **(A)** Fold- recognition analysis of KpsD*_Ec_* (via HHpred) revealing C-terminal structural homology to GfcC (PDB: 3P42) as well as Wza*_Ec_* (PDB: 2J58). **(B)** Profile-based alignment of Wza*_Ec_* and KpsD*_Ec_* C-terminal sequences from Panel A. Wza*_Ec_* α-helix (*dark green cylinders*) and β-strand (*dark blue arrows*) structure is depicted as per the 2J58 PDB entry. KpsD*_Ec_* predicted α-helix (*light green cylinders*) and β-strand (*light blue arrows*) secondary structure is indicated as per PSIPRED analysis. Aligned residues have been coloured according to Jalview conservation score (out of 10). *Maroon*, 10; *red*, 9; *orange*, 8; *yellow*, 7; *pink*, 6. Scores of 5 or less have been omitted to improve clarity of the figure. The end of KpsD*_Ec_* structural homology with the stand-alone GfcC protein has been indicated as per a previous report (Sande *et al*., 2019). **(C)** Tertiary structure model of the KpsD*_Ec_* C-terminus based on structural alignment with Wza*_Ec_* as indicated in Panel B. N- and C-termini of the displayed peptide have been indicated.

**Supplementary Figure S2.**
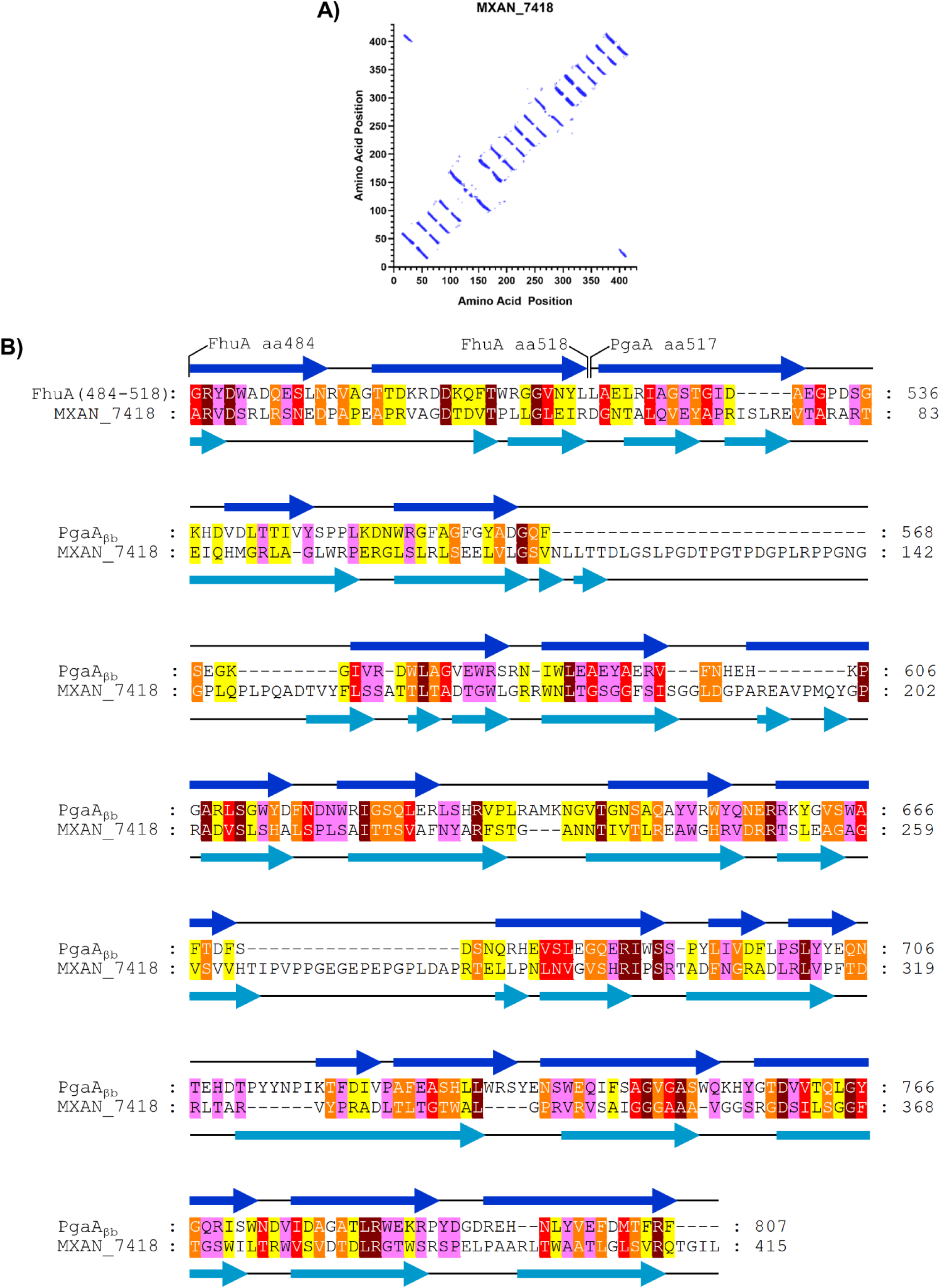
Structural analysis of MXAN_7418 (WzpX). **(A)** Evolutionarily- coupled amino acids within the MXAN_7418 primary structure (determined via RaptorX). **(B)** Fold- recognition analysis of MXAN_7418 (via HHpred) revealing N-terminal structural homology with two β-strands from FhuA (PDB: 4CU4) (Mathavan *et al*., 2014), with the remainder of the protein displaying structural homology to PgaA_βb_ (PDB: 4Y25) (Wang *et al*., 2016). FhuA and PgaA_βb_ β- strand (*dark blue arrows*) structure is depicted as per the respective PDB entries. MXAN_7418 predicted β-strand (*light blue arrows*) secondary structure is indicated as per PSIPRED analysis. Aligned residues have been coloured according to Jalview conservation score (out of 10). *Maroon*, 10; *red*, 9; *orange*, 8; *yellow*, 7; *pink*, 6. Scores of 5 or less have been omitted to improve clarity of the figure.

**Supplementary Figure S3.**
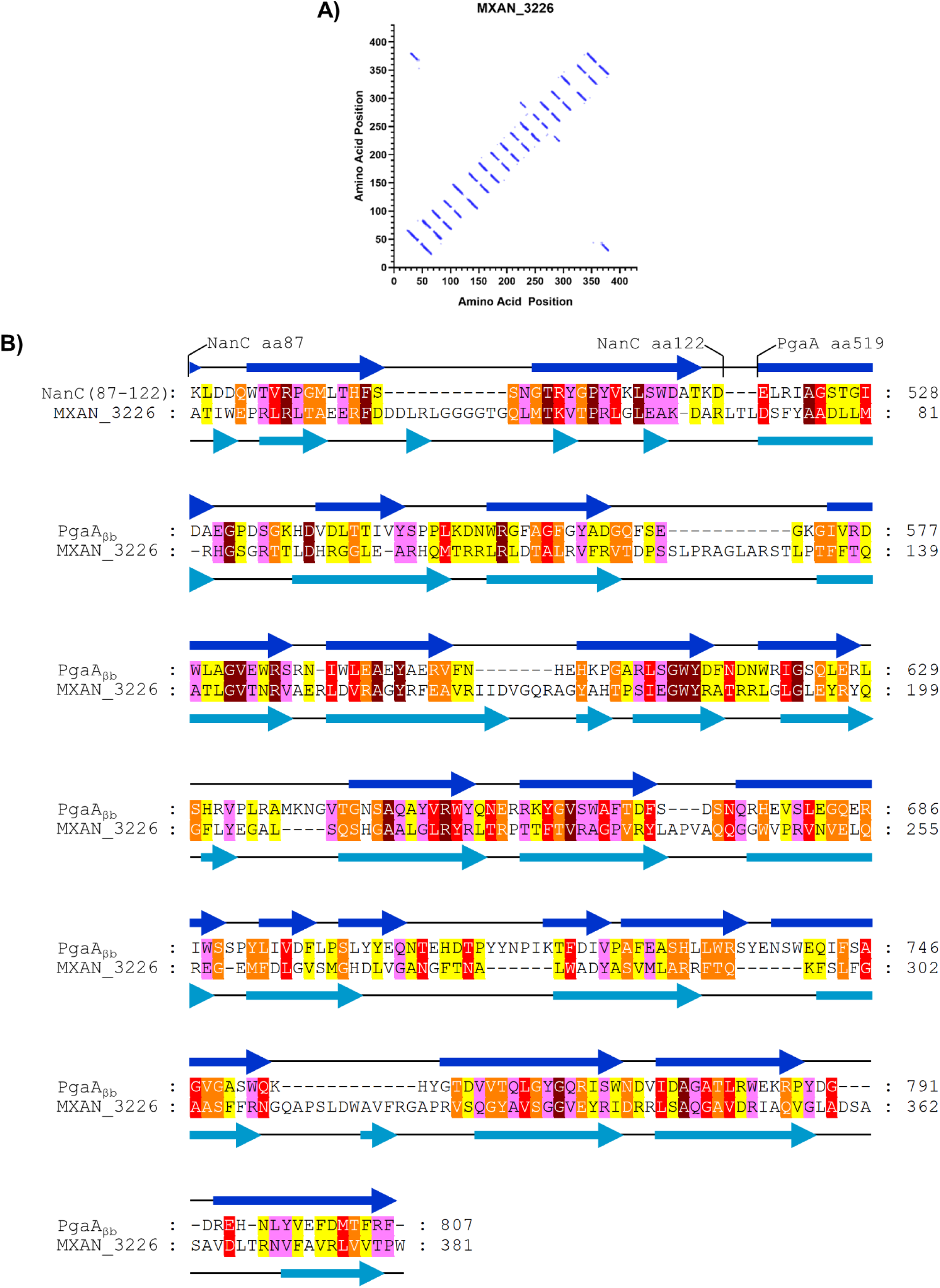
Structural analysis of MXAN_3226 (WzpS). **(A)** Evolutionarily- coupled amino acids within the MXAN_3226 primary structure (determined via RaptorX). **(B)** Fold- recognition analysis of MXAN_3226 (via HHpred) revealing N-terminal structural homology with two β-strands from NanC (PDB: 2WJR) (Wirth *et al*., 2009), with the remainder of the protein displaying structural homology to PgaA_βb_ (PDB: 4Y25) (Wang *et al*., 2016). NanC and PgaA_βb_ β-strand (*dark blue arrows*) structure is depicted as per the respective PDB entries. MXAN_3226 predicted β- strand (*light blue arrows*) secondary structure is indicated as per PSIPRED analysis. Aligned residues have been coloured according to Jalview conservation score (out of 10). *Maroon*, 10; *red*, 9; *orange*, 8; *yellow*, 7; *pink*, 6. Scores of 5 or less have been omitted to improve clarity of the figure.

**Supplementary Figure S4.**
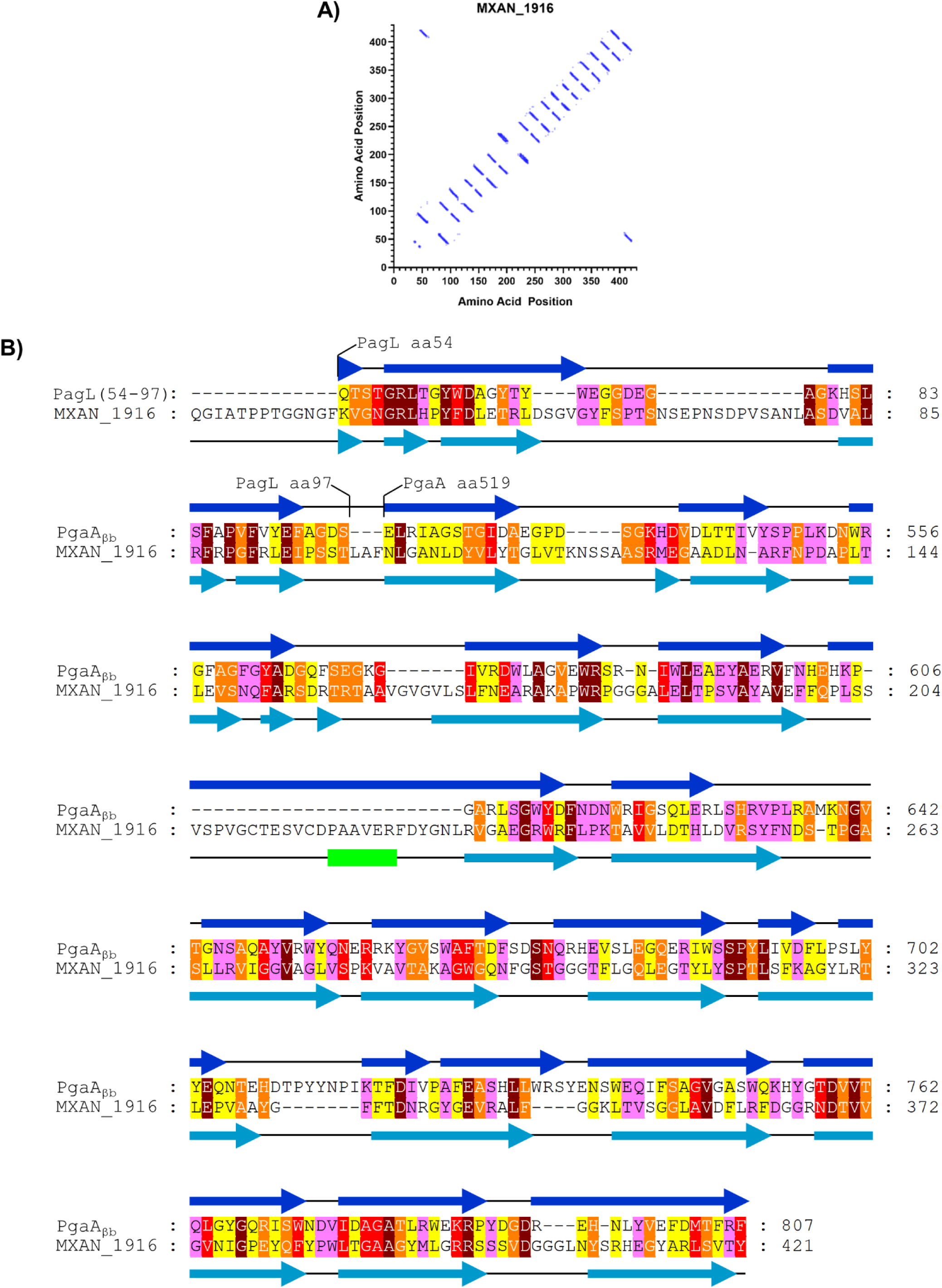
Structural analysis of MXAN_1916 (WzpB). **(A)** Evolutionarily- coupled amino acids within the MXAN_1916 primary structure (determined via RaptorX). **(B)** Fold- recognition analysis of MXAN_1916 (via HHpred) revealing N-terminal structural homology with two β-strands from PagL (PDB: 2ERV) (Rutten *et al*., 2006), with the remainder of the protein displaying structural homology to PgaA_βb_ (PDB: 4Y25) (Wang *et al*., 2016). PagL and PgaA_βb_ β-strand (*dark blue arrows*) structure is depicted as per the respective PDB entries. MXAN_1916 predicted α-helix (*light green cylinders*) and β-strand (*light blue arrows*) secondary structure is indicated as per PSIPRED analysis. Aligned residues have been coloured according to Jalview conservation score (out of 10). *Maroon*, 10; *red*, 9; *orange*, 8; *yellow*, 7; *pink*, 6. Scores of 5 or less have been omitted to improve clarity of the figure.

**Supplementary Figure S5.**
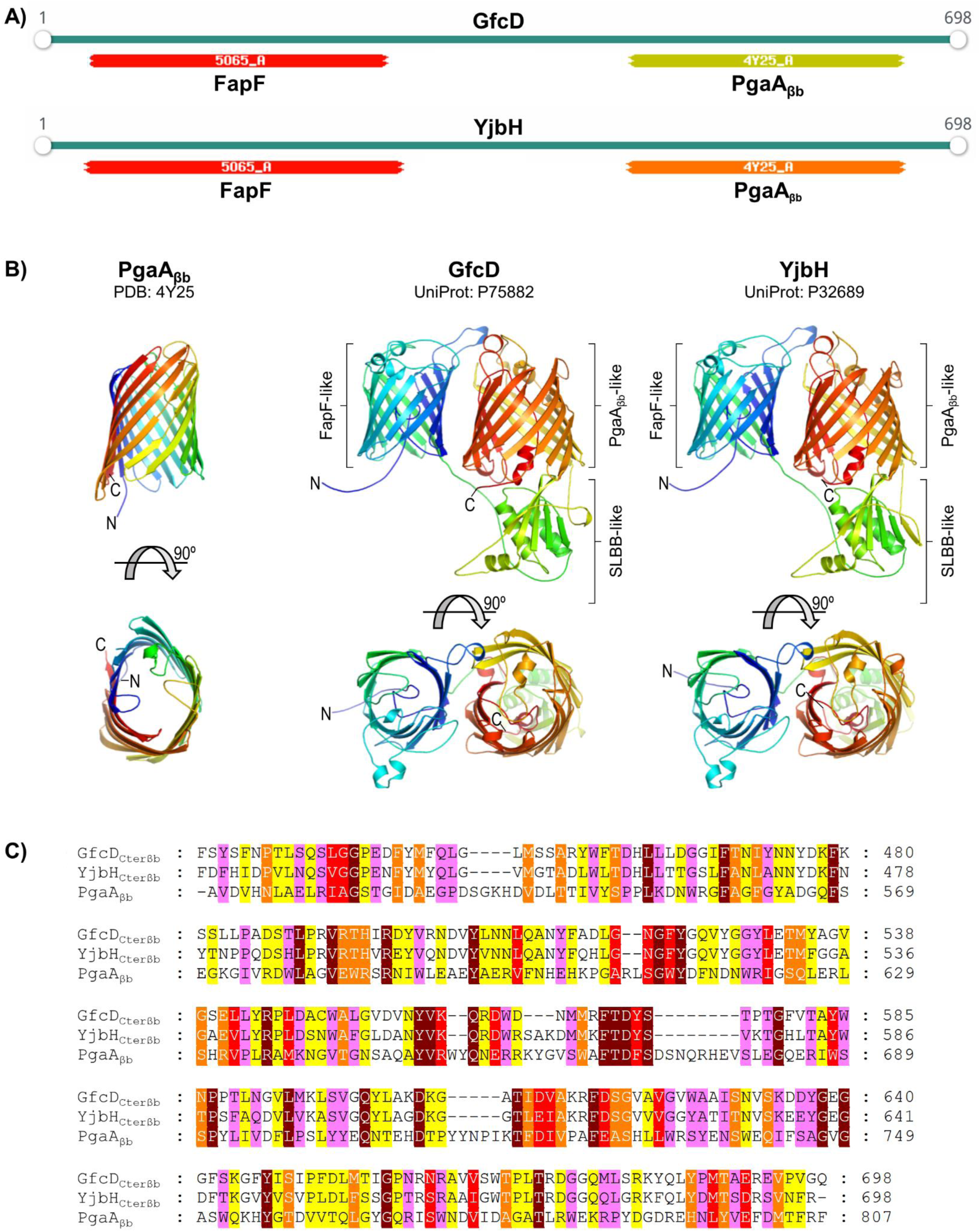
Structural homology between Pgaβb, GfcD, and YjbH. **(A)** Fold- recognition analysis of GfcD and YjbH (via HHpred) revealing N-terminal structural homology of each to the amyloid secretion β-barrel FapF (PDB: 5O65) (Rouse *et al*., 2017) and C-terminal structural homology of each to PgaA_βb_ (PDB: 4Y25) (Wang *et al*., 2016). **(B)** AlphaFold2-generated tertiary structure models for GfcD and YjbH, displayed alongside the Pga_βb_ X-ray crystal structure for comparison. Proteins have been coloured with a spectrum, from the N-terminus (*blue*) to the C- terminus (*red*). **(C)** Multiple-sequence alignment of the PgaA_βb_ (aa 511-807), GfcD_Cterβb_ (aa 425- 698), and YjbH_Cterβb_ (aa 423-698) segments. Aligned residues have been coloured according to Jalview conservation score (out of 10). *Maroon*, 10; *red*, 9; *orange*, 8; *yellow*, 7; *pink*, 6. Scores of 5 or less have been omitted to improve clarity of the figure.

**Supplementary Figure S6.**
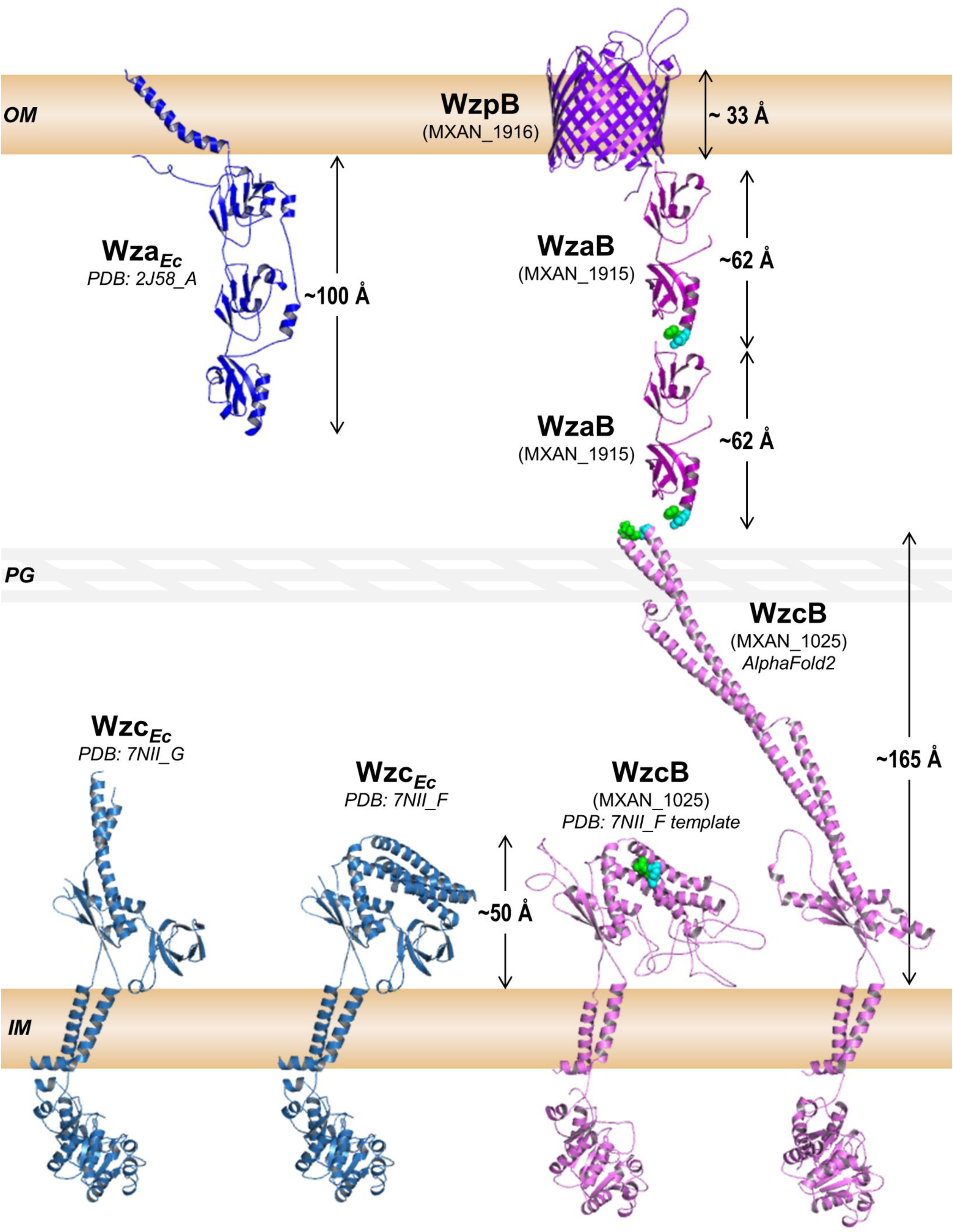
Structural schematic of polymer translocation across the periplasm in *M. xanthus*. Components from the BPS secretion pathway have been used as representative proteins for those in the EPS and MASC pathways as well. All proteins, spaces, and distances have been depicted at the same relative scale across a representative 327 Å periplasmic space in a *M. xanthus* cell. X-ray crystal structures for Wza*_Ec_* (chain A) and Wzc*_Ec_* (chains F and G) have been provided as per the PDB files 2J58 (Dong *et al*., 2006) and 7NII (Yang *et al*., 2021) (respectively) for size references. Structure models for WzaB (Fig. 1B) and WzpB (Fig. 5A) were already generated in this investigation. Models for the PCP protein WzcB were generated using either AlphaFold2 (resulting in an extended conformation), or MODELLER (specifically against the 7NII_F template, resulting in a compact conformation). High-confidence co-evolving amino acids between WzcB and WzaB have been highlighted with *green* (86% probability) and *cyan* (81% probability) spheres.

## SUPPLEMENTARY TABLE LEGENDS

**Supplementary Table S1. MYXO dataset analysis. (A)** Protein-wise OPX classification within 61 myxobacterial genomes. **(B)** Distribution of OPX-protein types within myxobacterial genomes. **(C)** Synteny analysis of OPX-protein types with β-barrel protein homologues.

**Supplementary Table S2. REP dataset analysis. (A)** Protein-wise OPX classification within 3662 Representative/Reference genomes. **(B)** Distribution of OPX-protein Classes within 3662 genomes. **(C)** Synteny distribution of OPX-protein types with β-barrel protein homologues. **(D)** Arrangement of syntenic OPX-protein types with β-barrel protein homologues.

**Supplementary Table S3. NR dataset analysis. (A)** Protein-wise OPX classification within NR- database proteins.

**Supplementary Table S4. Distribution of OPX-protein types at Phylum, Class, Order, Family, and Genus level taxonomy.**

**Supplementary Table S5. Evolutionary couplings between proteins.** Co-evolving amino acids between PCP and OPX pairs, as well as OPX and Wzp β-barrel pairs, are presented for constituents of the EPS, BPS, and MASC pathways.

